# PfAP2-MRP DNA-binding protein is a master regulator of parasite pathogenesis during malaria parasite blood stages

**DOI:** 10.1101/2023.05.23.541898

**Authors:** Amit Kumar Subudhi, Judith L. Green, Rohit Satyam, Todd Lenz, Rahul P. Salunke, Muhammad Shuaib, Ioannis Isaioglou, Steven Abel, Mohit Gupta, Luke Esau, Tobias Mourier, Raushan Nugmanova, Sara Mfarrej, Rupali Sivapurkar, Zenaida Stead, Fathia Ben Rached, Yogesh Otswal, Rachid Sougrat, Ashraf Dada, Abdullah Fuaad Kadamany, Wolfgang Fischle, Jasmeen Merzaban, Ellen Knuepfer, David J.P. Ferguson, Ishaan Gupta, Karine G. Le Roch, Anthony A. Holder, Arnab Pain

## Abstract

Malaria pathogenicity results from the parasite’s ability to invade, multiply within and then egress from the host red blood cell (RBC). Infected RBCs are remodeled, expressing antigenic variant proteins (such as PfEMP1, coded by the *var* gene family) for immune evasion and survival. These processes require the concerted actions of many proteins, but the molecular regulation is poorly understood. We have characterized an essential *Plasmodium* specific Apicomplexan AP2 (ApiAP2) transcription factor in *Plasmodium falciparum* (PfAP2-MRP; Master Regulator of Pathogenesis) during the intraerythrocytic developmental cycle (IDC). An inducible gene knockout approach showed that PfAP2-MRP is essential for development during the trophozoite stage, and critical for *var* gene regulation, merozoite development and parasite egress. ChIP-seq experiments performed at 16 hour post invasion (h.p.i.) and 40 h.p.i. matching the two peaks of PfAP2-MRP expression, demonstrate binding of PfAP2-MRP to the promoters of genes controlling trophozoite development and host cell remodeling at 16 h.p.i. and antigenic variation and pathogenicity at 40 h.p.i. Using single-cell RNA-seq and fluorescence-activated cell sorting, we show de-repression of most *var* genes in *Δpfap2-mrp* parasites that express multiple PfEMP1 proteins on the surface of infected RBCs. In addition, the *Δpfap2-mrp* parasites overexpress several early gametocyte marker genes at both 16 and 40 h.p.i., indicating a regulatory role in the sexual stage conversion. Using the Chromosomes Conformation Capture experiment (Hi-C), we demonstrate that deletion of PfAP2-MRP results in significant reduction of both intra-chromosomal and inter-chromosomal interactions in heterochromatin clusters. We conclude that PfAP2-MRP is a vital upstream transcriptional regulator controlling essential processes in two distinct developmental stages during the IDC that include parasite growth, chromatin structure and *var* gene expression.

## Main

Malaria pathology is caused by asexual blood stage parasite growth. Merozoite invasion of red blood cells (RBCs), intraerythrocytic replication and then egress from infected RBC (iRBCs) drive proliferation^1^. The parasite modifies the iRBC surface, displaying antigenically variant proteins encoded by multicopy gene families to escape immune clearance^2^. These processes involve concerted expression of hundreds of proteins during specific periods of the intraerythrocytic development cycle (IDC)^1–3^, including proteins encoded by genes that show clonal variant expression^4^. Regulation of gene expression involves sequence-specific DNA binding proteins as transcriptional activators or repressors and epigenetic modifiers^5^. Chromatin-mediated regulation of *P. falciparum* gene expression controls invasion proteins and antigenically variant proteins^4^. Recruitment of epigenetic regulators is poorly understood. A likely possibility is that they are recruited via DNA-binding regulatory proteins. In the genus *Plasmodium*, a family of 27 Apicomplexan-specific ApiAP2 DNA-binding proteins has been identified as the major transcriptional regulators of various processes during development, differentiation, and response to environmental changes^6–11^. However, few have been characterized in detail to decipher their function and the role of the remainder is unknown.

Here, we use an inducible gene knockout approach^12, 13^ to show that PF3D7_1107800 codes for an ApiAP2 (PfAP2-MRP) essential for cell cycle, antigenic variation and parasite egress and invasion. We establish that PfAP2-MRP directly or indirectly regulates the expression of genes involved in host cell remodeling, antigenic variation, egress, invasion and gametocytogenesis amongst the pathogenesis-related processes corresponding to the two peaks of expression during the IDC. We also provide ChIP-seq evidence that PfAP2-MRP binds to the promoter regions of genes predicted to be involved in transcription, translation and nucleosome assembly. Chromosomes Conformation Capture (Hi-C) analysis demonstrates that deletion of PfAP2-MRP results in significant reduction of heterochromatin clusters leading to transcriptional activation of gene involved in antigenic regulation and sexual differentiation. We conclude that PfAP2-MRP is a master regulator of parasite growth, chromatin structure and *var* genes expression.

### PfAP2-MRP is essential for parasite development and growth

In a recent study^14^, we identified 363 genes with transcripts displaying a 24 h (circadian-like) rhythmic periodicity in the 48-hour *P. falciparum* IDC, one of which was an ApiAP2 we now designate PfAP2-MRP (Extended Data Fig. 1a). The first peak of expression at ∼16 h.p.i. coincides with maximal *var* gene family expression. The second peak at ∼ 40 h.p.i., is just before maximal expression of known invasion- and egress-associated genes, suggesting a role of PfAP2-MRP in regulating these genes (Extended Data Fig. 1a).

PfAP2-MRP has one AP2 domain (residues 1487-1544, PFAM ID:PF00847), encoded in the second exon (Extended Data Fig. 1b). PfAP2-MRP orthologs are found only in *Plasmodium* spp. and in no other members of Apicomplexa (Extended Data Fig. 1c). The PfAP2-MRP and its *P. berghei* ortholog (PBANKA_0939100) were identified as essential during blood-stage development^15, 16^. To investigate the function of PfAP2-MRP, we generated a 3HA-tagged PfAP2-MRP inducible knockout *P. falciparum* (PfAP2-MRP-3HA:loxP) using a parental line that expresses rapamycin (RAPA)-inducible dimerizable Cre recombinase (DiCre)^13^ (Fig. 1a, Methods).

**Fig. 1.**
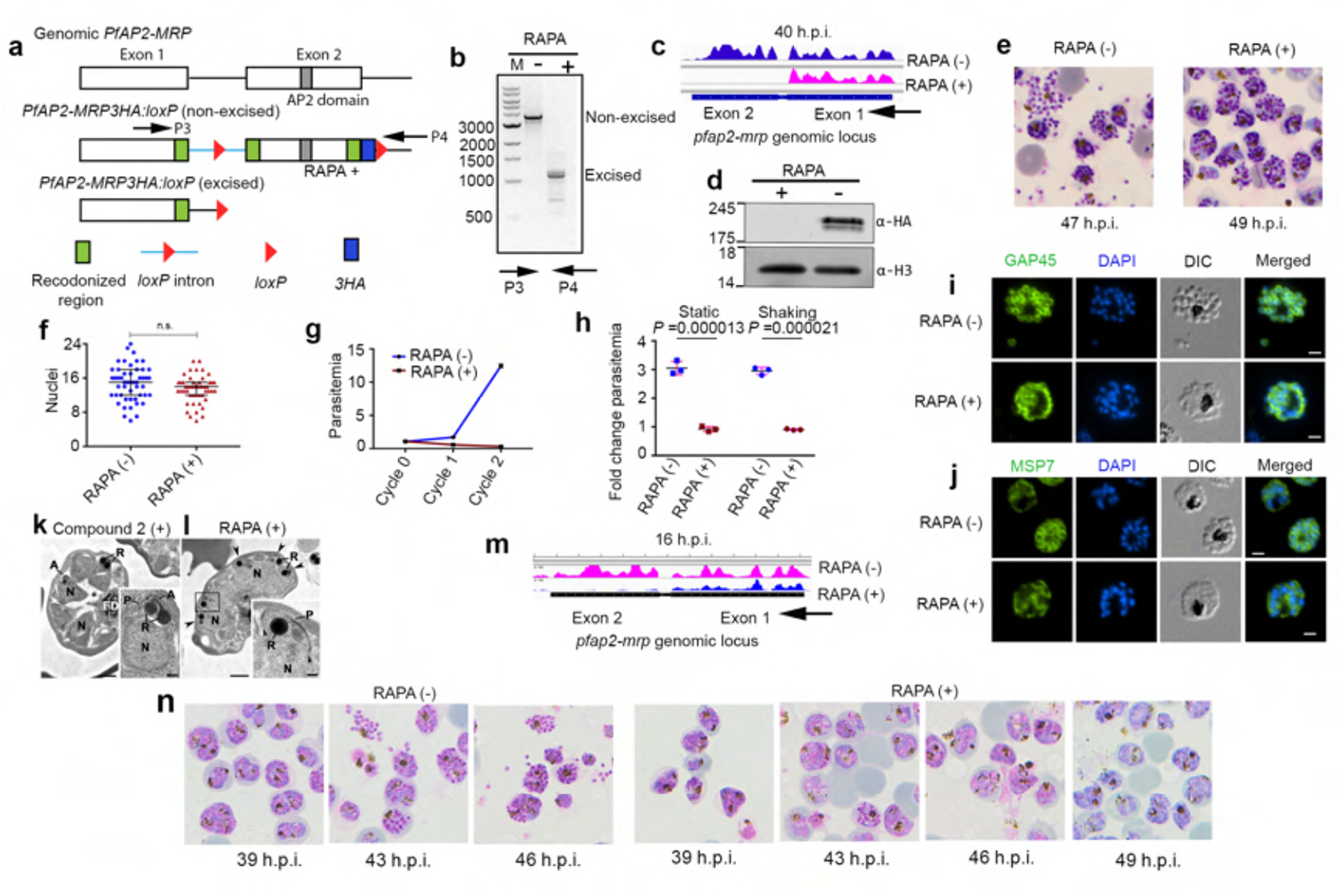
PfAP2-MRP is essential for parasite growth and development. **a,** Schematic of strategy to tag *pfap2-mrp* with 3HA sequence and its conditional knockout by excision of the *loxP-*flanked second exon. **b**, Diagnostic PCR confirms exon2 excision in RAPA-treated parasites. **c,** RNA-seq reads coverage from 40 h.p.i. parasites of *pf-ap2-mrp* locus. **d,** Western blot of control (RAPA -) and rapamycin (RAPA +) treated PfAP2-MRP-3HA:*loxP* schizont extracts probed with anti-HA and -histone H3 antibodies. Molecular mass (kDa) of standards on left side of each panel. **e,** Images of Giemsa-stained (control [RAPA (-)] and treated [RAPA (+)]; treatment at 16 h.p.i) schizonts at end of cycle 0 (representative of 4 independent experiments). **f**, Nuclei in control [RAPA (-)] and treated [RAPA (+)] schizonts. The mean number was 14.7 (control) and 13.4 (treated); *P*-value of 0.178, one-way ANOVA using Sidak’s multiple comparisons test, assumed Gaussian distribution. **g**, Replication of control [RAPA (-)] and treated [RAPA (+)] parasites over two growth cycles (three biological replicates, average parasitemia ± s.d.). **h**, Erythrocyte invasion by control [RAPA (-)] and treated [RAPA (+)] PfAP2-MRP-3HA:*loxP* parasites under static and shaking conditions. Statistical significance: two-tailed t-test; RAPA (–) versus RAPA (+) parasites in static conditions (n=3, t = 14.17, d.f. = 4, P = 0.000013, 95% CI 1.606 to 2.389) and RAPA (–) versus RAPA (+) parasites in shaking conditions (n=3, t = 12.41, d.f. = 4, P = 0.000021, 95% CI 1.578 to 2.487). **i-j**, Immunofluorescence microscopy of control [RAPA (–)] and treated [RAPA (+)] parasites incubated with anti-GAP45 IgG (**i**) and anti-MSP7 serum (**j**); scale bar is 2 µm. **k-l**, Electron micrographs of iRBCs treated with compound 2(**k**) or rapamycin [RAPA(+)] (**l**); scale bar is 1 µm or 100 nm (inserts). **m**, RNA-seq reads coverage from 16 h.p.i. parasites (cycle 1) of *pf-ap2-mrp* locus. **n**, Images of Giemsa-stained parasites at 39, 43, 46 and 49 h.p.i. stages in cycle 1 following parasite treatment with DMSO [RAPA (-)] or rapamycin [RAPA(+)] at 35 h.p.i. at cycle 0.

Upon addition of RAPA to synchronized parasite culture, the *loxP*-flanked *pfap2-mrp* second exon was efficiently excised (Fig. 1b). A time-series RNA-seq analysis revealed that *loxP* flanked exon 2 deletion, and disruption of *pfap2-mrp* expression, occurred 14 to 16 hours after RAPA addition (Extended Data Fig. 1d), allowing us to determine when to add RAPA to disrupt each peak of *pfap2-mrp* expression during the IDC. To ablate only the second peak of functional *pfap2-mrp* expression at 40 h.p.i. in the cycle 0, RAPA was added at ∼16 h.p.i in the same cycle (i.e. cycle 0). To ablate only the first peak of expression at 16 h.p.i. in cycle 1, RAPA was added at ∼35 h.p.i. in cycle 0, with parasite collection at 16 h.p.i. in cycle 1 (Extended Data Fig. 1e). From here onwards, *Δpfap2-mrp* parasites (*Pfap2-mrp*-exon2 deleted) at 40 h.p.i. refers to parasites with disrupted second peak of expression that were collected at 40 h.p.i. and *Δpfap2-mrp* parasites at 16 h.p.i. refers to parasites with disrupted first peak of expression that were collected at 16 h.p.i.

RAPA addition at ∼16 h.p.i in cycle 0 ablated the second peak in the same cycle as demonstrated by RNA-seq (Fig. 1c) and western blot (Fig. 1d), and these *Δpfap2-mrp* parasites progressed through the IDC, forming morphologically normal segmented schizonts (49 h.p.i [cycle 0], Fig. 1e), with no significant difference in the number of nuclei in late schizonts of *Δpfap2-mrp* and DMSO (mock)-treated PfAP2-MRP-3HA:loxP parasites (Fig. 1f). However, *Δpfap2-mrp* parasites failed to egress (Fig. 1e), with a resultant dramatic reduction in parasitemia in subsequent cycles compared to controls (Fig. 1g), indicating an essential role of PfAP2-MRP in parasite proliferation. In egress assays with highly synchronized segmented mature schizonts under either static or vigorous shaking (shear) conditions, no egress or increased parasitemia was observed for *Δpfap2-mrp* parasites, in contrast to the increased parasitemia in control cultures, irrespective of conditions (Fig. 1h); therefore, the blocked egress phenotype was not corrected by mechanical disruption of the iRBC.

To look for morphological differences, we compared *Δpfap2-mrp* and control parasites by indirect immunofluorescence with antibodies specific for proteins of the parasite surface pellicle: GAP45, a protein of the glideosome/inner membrane complex (IMC), and merozoite surface protein 7 (MSP7) (Fig. 1i and 1j). In *Δpfap2-mrp* compared with control parasites, the distribution of these proteins was disordered. Electron microscopy analysis of *Δpfap2-mrp* and parasites treated with compound 2 (an inhibitor of egress^17^) at 49 h.p.i. revealed that most mature schizonts contained fully formed merozoites in the control sample (Fig. 1k and Fig. 1k, insert). In contrast, most *Δpfap2-mrp* schizonts were at an earlier segmented stage (Fig. 1l and Fig. 1l, insert). In samples of 50 iRBCs, the control group contained 83% mature schizonts compared to the *Δpfap2-mrp* group that contained only 2% mature schizonts (Extended Data Fig. 1f). These data suggest that the function of the second peak of *pfap2-mrp* expression is manifest in the final IDC stages after nuclear division.

RAPA treatment from ∼35 h.p.i. had little effect on parasite egress or invasion at the end of cycle 0, but functional deletion of *pfap2-mrp* occurred well before the first expression peak in the next cycle. These parasites with the first peak of *pfap2-mrp* expression ablated were collected at 16 h.p.i. in cycle 1 (Extended Data Fig. 1e), and loss of functional *pfap2-mrp* expression was confirmed by RNA-seq (Fig. 1m). Parasite development stalled at the late trophozoite/early schizont stage in cycle 1 (Fig. 1n), suggesting that the first peak of *pfap2-mrp* expression during the IDC plays a critical role in parasite development immediately after its expression at ∼16 h.p.i.

### PfAP2-MRP is a crucial regulator of malaria pathogenesis associated genes

A comparative RNA-seq analysis of *Δpfap2-mrp* and control parasite populations at 16 and 40 h.p.i. identified 793 and 1,389 differentially expressed genes (FDR ≤ 0.05), respectively (Extended Data Fig. 2a,b and Supplementary Data 1). Because disruption of the second peak of *pfap2-mrp* expression caused parasites to stall at the late segmented stage, we examined whether the observed transcriptional differences at 40 h.p.i. were due to the knockout or reduced viability. We focused on the expression of 1,042 genes that are known to have peak expression after 35 h.p.i.^5^, and expected that death or a delay in development of *Δpfap2-mrp* parasites would result in most of these genes being identified as down-regulated. However, 658 (63%) of them showed no significant change in expression level (Extended Data Fig. 2c), indicating that observed differences in gene expression were due to *pfap2-mrp* deletion and not reduced viability.

The *P. falciparum* genome contains 59 intact *var* genes and their expression is mutually exclusive: a single parasite expresses one *var* gene at a time, with the remainder of the family remaining transcriptionally silent by heterochromatin formation^18^. Gene ontology (GO) enrichment analysis of up-regulated genes in 16 and 40 h.p.i *Δpfap2-mrp* parasites identified pathogenesis as the most enriched biological term (adjusted *P* = 1.23e-07 and 6.8e-10, respectively; Supplementary Data 2). The *var* gene family has the largest number of members assigned to this GO term, with significant up-regulation of 24 and 29 *var* genes (*Padj* < 0.05) in *Δpfap2-mrp* compared to control parasites at 16 h.p.i. and 40 h.p.i., respectively (Fig. 2a,b). Data was validated by quantitative real time PCR (qRT-PCR) (Fig. 2c and 2d). These results suggest that PfAP2-MRP may acts a repressor of most *var* genes.

**Fig. 2.**
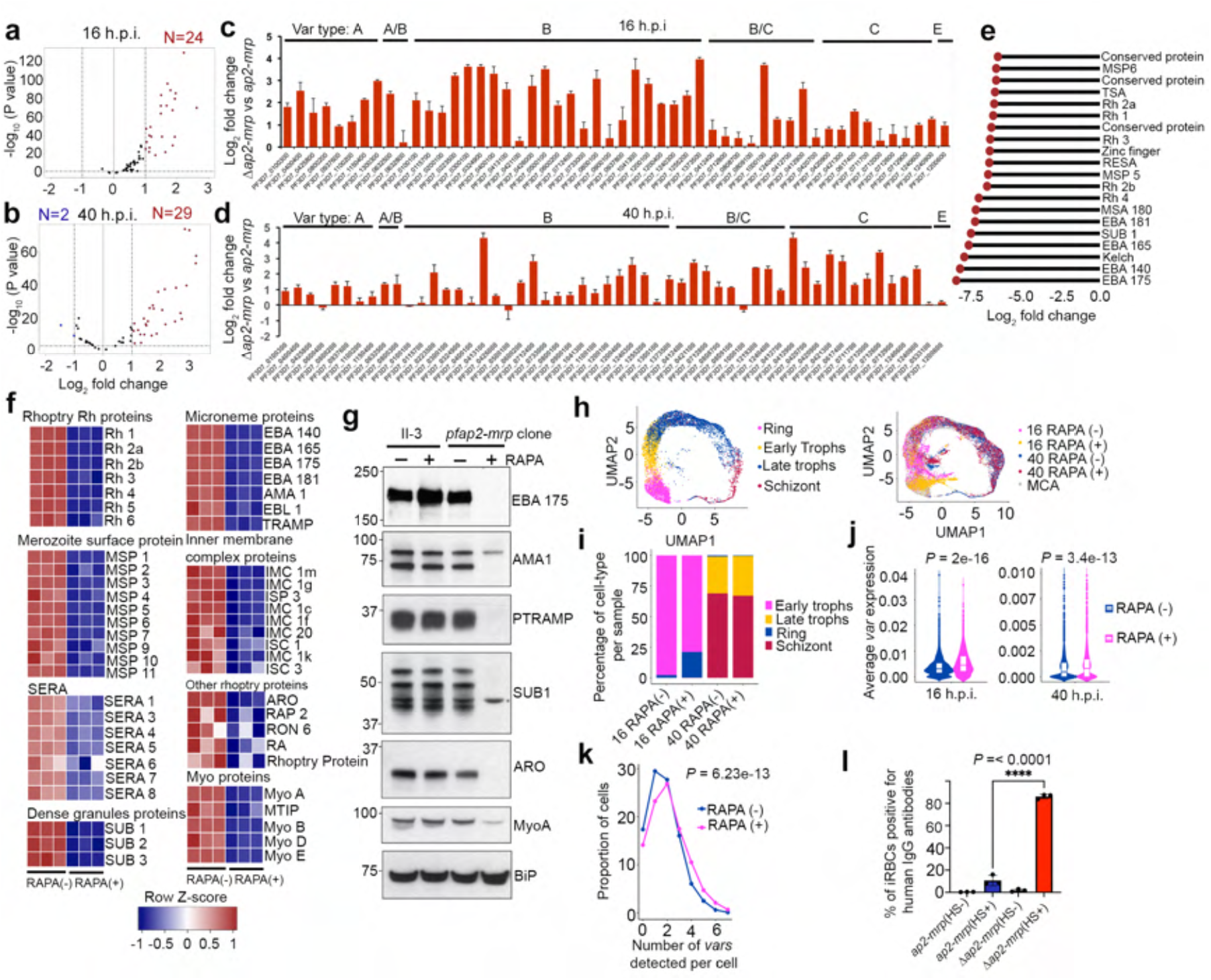
PfAP2-MRP regulates most of the malaria pathogenesis-associated genes. **a**-**b**, Volcano plots showing differentially expressed *var* genes in 16 h.p.i (cycle 1) (**a**) and 40 h.p.i. (cycle 0) (**b**) *Δpfap2-mrp* parasites. **c-d**, Differential expression of *var* genes (log2 ratio of Δ*pfap2-mrp* to mock-treated control parasites) measured by qRTPCR at 16 h.p.i. (**c**) and 40 h.p.i. (**d**); error bar is s.e.m. **e**, Lollipop plot of expression level of top 20 significantly down-regulated genes in Δ*pfap2-mrp* compared to control parasites at 40 h.p.i. **f,** Heatmaps for most down-regulated genes known to be involved in parasite egress and invasion in treated [RAPA (+)] or control [RAPA (-)] parasites, grouped based on the sub-cellular location of their products. **g**, Western blots of schizont extract from parental II-3 and PfAP2-MRP-3HA parasites in the absence (-) or presence (+) of rapamycin, probed with antibodies specific for invasion proteins. BiP was detected as a loading control. Molecular mass (kDa) of standards on left side of each panel. A non-specific cross-reacting protein on the SUB1 blot is marked with an asterisk. **h**, Left: uniform manifold approximation and projection (UMAP) of scRNA-seq data from Malaria Cell Atlas (MCA), with annotated developmental stages. Right: UMAP projections of scRNA-seq in-house data; each dot represents gene expression data from a single parasite (colours corresponding to h.p.i. and *pfap2-mrp* knockout status) plotted over MCA data. **i**, Distribution of developmental stages of treated [RAPA(+)] or control [RAPA(-)] parasites at 16 and 40 h.p.i. **j**, Violin plots of average *var* gene expression in treated [RAPA (+)] or control [RAPA (-)] parasites at both 16 h.p.i. and 40 h.p.i. **k**, Proportion of cells expressing one or more *var* genes from treated [RAPA(+)] or control [RAPA (-)] cultures (*P*= 6.23e-13, Fisher’s exact test, odds ratio=1.53). **l**, Percentage of iRBCs containing control or Δ*pfap2-mrp* parasites bound by IgG from serum of malaria-infected (HS+) patients, or untreated samples (HS-) . Significance determined using a two-tailed t-test (t = 6.687, d.f. = 4, *P* < 0.0001, 95% CI 0.9844 to 2.382, n=3).

We also observed up-regulation of 4 of 8 *surfins* at 16 h.p.i. (Supplementary Data 1). Several gene families coding for other variant proteins such as *rifins* (n=78/132), *stevors* (n=13/30), and *Pfmc-2ms* (n=13/13) were significantly down-regulated (FDR ≤ 0.05) at 16 h.p.i., suggesting that these genes are positively regulated by PfAP2-MRP (Extended Data Fig. 2d, Supplementary Data 1). While some members of the *hyp* and *phist* gene families, encoding exportome-associated proteins showed up-regulation, others showed down-regulation at 16 h.p.i., consistent with a functional diversification (Supplementary Data 1). Together, these results suggest that *pfap2-mrp* expression at 16 h.p.i. has a major role in regulating the expression of genes coding for proteins important in antigenic variation and host cell remodeling.

Gene ontology (GO) enrichment analysis of the down-regulated genes in *Δpfap2-mrp* parasites at 40 h.p.i. identified entry into and egress from the host cell as the two most enriched biological process terms (Extended Data Fig. 2e, Supplementary Data 2), consistent with our finding that deletion of *pfap2-mrp* critically affects late merozoite development and egress. Of the top twenty down-regulated genes, fifteen function in parasite egress or invasion (Fig. 2e). The expression of all top down-regulated genes peaks just after the second peak of *pfap2-mrp* expression (Extended Data Fig. 1a), consistent with the proposed role of PfAP2-MRP in controlling their expression. For example, known egress-associated genes coding for SUB1 -2 and -3, SERA5 and -6, Plasmepsin X, CDPK1 and -5, MSA 180, phospholipase, PKG, and MSP1 were down-regulated (Supplementary Data 1). Strikingly, 36 out of 72 known invasion-associated genes, including those coding for micronemal, rhoptry, merozoite surface, IMC and motor proteins were also down-regulated (Fig. 2f, Supplementary Data 1). Western blots and immunofluorescence assays were used to confirm the reduced abundance for a few selected proteins (Fig. 2g). Altogether, the data suggest that PfAP2-MRP is essential to the expression of egress and invasion-associated genes.

Other biological processes enriched in the group of down-regulated genes in *Δpfap2-mrp* parasites at 40 h.p.i. include protein phosphorylation, actin cytoskeleton organization, and fatty acid elongation. Protein phosphorylation has a significant role in control of both egress and invasion in addition to other cell cycle events^19, 20^. We identified 30 out of 105 genes annotated as involved in protein phosphorylation to be significantly down-regulated (Extended Data Fig. 2f, Supplementary Data 2), including 10 out of 21 FIKK kinase genes, the majority of which (17) are exported^21, 22^. This suggests a broad regulatory role for PfAP2-MRP in signaling processes during egress and invasion (Supplementary Discussion).

### PfAP2-MRP is essential for trophozoite stage development

We tested whether the gene expression pattern observed in the bulk RNA-seq data is also observed at single cell resolution. Single-cell RNA sequencing (scRNA-seq) was performed using parasites from 16 h.p.i. (control parasites and parasites with the first peak of expression disrupted) and 40 h.p.i. (control parasites and parasites with only the second peak of expression disrupted) stages. Based on the stage-specific annotation of cells in the malaria cell atlas (MCA)^23^, 16 h.p.i. cells are ring/early trophozoite parasites and 40 h.p.i. cells are late trophozoite and schizont stage parasites (Fig. 2h). We observed that more single *Δpfap2-mrp* parasites with disrupted first peak of expression at 16 h.p.i. were classified as ring rather than trophozoite compared to controls (Fig. 2i). However, no difference in stage was observed at 40 h.p.i. for parasites with disrupted second peak of expression suggesting that it is not RAPA toxicity that is responsible for the observed delay of *Δpfap2-mrp* parasite development at 16 h.p.i. The delay observed in the parasite development may be an early phenotype resulting from the loss of the first *pfap2-mrp* expression peak (Fig. 1n), which suggest that the first peak of expression is critical for trophozoite development. No difference in developmental stages between control and *Δpfap2-mrp* parasites with only disrupted second peak of expression at 40 h.p.i. further suggests that differentially expressed genes at 40 h.p.i. (Extended Data Fig. 2b) were due to *pfap2-mrp* deletion and not due to difference in developmental stage or reduced viability.

### The role of PfAP2-MRP in repressing *var* genes confirmed with single cells

From the scRNA-seq analysis, we observed significantly higher expression of *var*s (Fig. 2j) and *surfins* (Extended Data Fig. 2g) and down-regulation of *rifins* and *pfmc-2tms* in *Δpfap2-mrp* parasites, supporting the bulk RNA-seq data (Extended Data Fig. 2g). We further examined whether *pfap2-mrp* deletion activated expression of multiple *var* genes in a single cell. Most control parasites expressed only one *var* gene following the mutually exclusive expression pattern reported for *var* genes^18^, but a few expressed two or more *var* genes (Fig. 2k). A significantly higher number of *Δpfap2-mrp* parasites expressed two or more *var* genes (*P*= 6.23e^-13^, Fisher’s exact test, Fig. 2k), indicating apparent disruption of mutually exclusive *var* gene expression in the *Δpfap2-mrp* parasites.

### *Δpfap2-mrp* parasites expresses multiple surface PfEMP1

To determine whether in *Δpfap2-mrp* parasites expressing multiple *var* genes, EMP1s are translated and exported to the iRBC surface, a FACS-based assay was developed with pooled serum from patients infected with *P. falciparum*^24^. Amongst many exported malaria proteins, PfEMP1 is a major target of naturally acquired antibodies^25^, and therefore we hypothesized that if *Δpfap2-mrp* parasites express multiple PfEMP1s, then they will bind more antibodies in the serum against different PfEMP1s, compared to the control parasites. We observed significantly more *Δpfap2-mrp* iRBCs than control parasites with bound antibody (*P*< 0.0001, two-tailed t-test, Fig. 2l, Extended Data Fig. 2h). This result indicates that activation of multiple *var* genes and the translation and transport of PfEMP1 to the iRBC surface occurs in *Δpfap2-mrp* parasites, and suggests that PfAP2-MRP is involved broadly in silencing *var* gene expression.

### PfAP2-MRP is a repressor of early gametocyte-associated marker genes

GO enrichment analysis of 526 up-regulated genes at 40 h.p.i. in *Δpfap2-mrp* schizonts identified significant enrichment of many biological processes, including lipid and fatty acid metabolism (Extended Data Fig. 3a and Supplementary Data 2). Genes such as *elongation of fatty acids protein 3* (*elo3*) and *acyl-CoA synthetase 9* (*acs9*) have essential roles in gametocyte development^16, 26^ and were up-regulated in 16 and 40 h.p.i. *Δpfap2-mrp* parasites (Supplementary Data 1). We examined the list of up-regulated genes for other known or putative early gametocyte marker genes^27, 28^ and found 18 out of 28 genes to be up-regulated in both 16 and 40 h.p.i. *Δpfap2-mrp* parasites (Extended Data Fig. 3b, Supplementary Data 1), suggesting that PfAP2-MRP is also a repressor of commitment to sexual stage development (Supplementary Discussion). We validated the expression of some differentially regulated genes by qRT-PCR, and found strong agreement between RNA-seq and qRT-PCR data (Extended Data Fig. 3c).

### PfAP2-MRP is a direct regulator of heterochromatin-associated genes

To distinguish between direct and indirect targets of PfAP2-MRP, we performed chromatin immunoprecipitation with the 3HA-epitope tagged PfAP2-MRP protein followed by sequencing (ChIP-seq). We identified 1,081 and 640 ChIP-seq peaks at 16 and 40 h.p.i., respectively, of which 78% and 68% were in intergenic/promoter regions upstream of at least one gene (with most located less than 2 kb upstream of an ATG translational start site) (Extended Data Fig. 4a and 4b, Supplementary Data 3). We validated a few PfAP2-MRP binding regions identified by ChIP-seq at 40 h.p.i. using ChIP-qPCR, with complete agreement (Extended Data Fig. 4c). Consistent with previous finding^29^, we observed binding of PfAP2-MRP to both central and sub-telomeric heterochromatin regions at both 16 and 40 h.p.i. stages in addition to its binding to the promoter of euchromatic genes (Extended Data Fig. 4d). GO analysis identified encoded proteins enriched in several processes, the most significant of which is antigenic variation (*Padj*=8.36e-32 at 16 h.p.i and 6.42e-38 at 40 h.p.i.; Supplementary Data 4). We observed that PfAP2-MRP binds significantly (*q* < 0.05) to the promoters of at least 37 and 33 *var* genes at 16 h.p.i. and 40 h.p.i, respectively; a total of 45 *var* genes, including both sub-telomeric and internal *var* genes of all the upstream sequence (ups) types (Fig. 3a and Extended Data Fig. 4e).

**Fig. 3.**
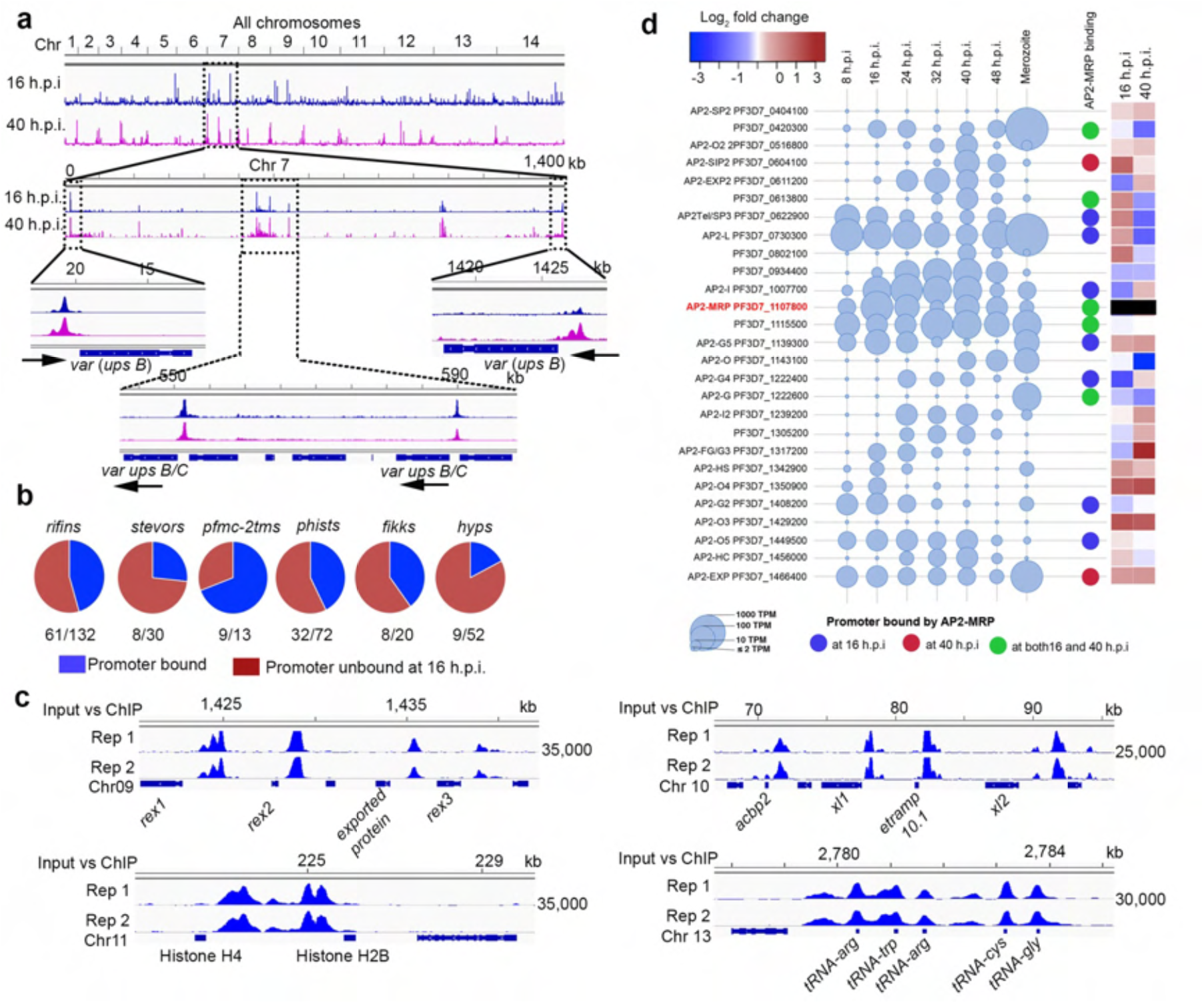
PfAP2-MRP regulates pathogenesis-associated genes via promoter binding. **a**, Genome-wide occupancy of PfAP2-MRP at 16 h.p.i. and 40 h.p.i., determined by ChIP-seq. Sub-telomeric and internal regions of chromosome 7 (∼1450 kb) containing *var* genes with PfAP2-MRP bound in their promoter regions are shown as an example. Chromosomal positions are indicated. Results are representative of 2 independent replicates. **b**, Pie charts showing the proportion of each family of genes with PfAP2-MRP bound to the promoter region (in blue) at 16 h.p.i. **c**, PfAP2-MRP occupancy at 16 h.p.i. in putative promoter regions of genes implicated in iRBC remodeling and parasite development. Two biological replicate ChIP vs. Input tracks are shown, (input subtracted PfAP2-MRP-ChIP). Positions on Chr 09, 10, 11 and 13 are indicated. X-axis shows the genomic position and numbers on the right, show the enrichment score (see Methods). **d**, Expression levels of 27 *P. falciparum* AP2 genes during different IDC stages and in merozoites are depicted by the diameter of the circles. On the right, blue, brown and green circles indicate the binding of PfAP2-MRP to the promoter at either or both 16 h.p.i. and 40 h.p.i.; the heatmap displays the expression status of all 26 *api-ap2s* in Δ*pfap2-mrp* parasites compared to controls at 16 and 40 h.p.i. The heatmap for *pfap2-mrp* is black because *Δpfap2-mrp* parasites only express RNA from the first exon and have no functional AP2-MRP protein.

PfAP2-MRP binding at both 16 and 40 h.p.i. was also enriched at the promoter of genes encoding exportome proteins with relatively strong signals at 16 h.p.i stage and at the promoter or intragenic regions of many other gene families for antigenic variant proteins (*rifins*, *stevor*, *surfins* and *pfmc-2tms*) with relatively weaker yet statistically significant signals at 16 h.p.i. stage (Fig. 3b and 3c, Supplementary Data 3). Most of these genes were also differentially expressed in *Δpfap2-mrp* parasites suggesting that they are under direct PfAP2-MRP control (Supplementary Data 1). These gene families for antigenic variant and other exported proteins are located within the chromosomal central and subtelomeric heterochromatin regions.

The promoters of genes associated with critical processes in early-stage development such as translation (*Padj*=1.3e-10), glycolytic process (*Padj*=0.0005) and nucleosome assembly (*Padj*= 0.02) were also strongly bound by PfAP2-MRP at 16 h.p.i. (Fig. 3c, Supplementary Data 3 and 4) suggesting these processes are regulated by PfAP2-MRP, and to some extent this may explain why development of parasites with disrupted first peak of expression stalled at late trophozoite stage.

### PfAP2-MRP binds to its own promoter and those of another 13 *ApiAP2*s

PfAP2-MRP binds to the promoter of a total of 14 *Pfapiap2* genes including its own promoter at both 16h.p.i. and 40 h.p.i. (Fig. 3d, Extended Data Fig. 5 and Supplementary Data 3). Nine of these 13 *apiap2s* were differentially expressed in *Δpfap2-mrp* parasites at one or both of the 16 and 40 h.p.i. time points, suggesting that they are under the direct control of PfAP2-MRP.

We observed binding of PfAP2-MRP to the *pfap2-i* promoter (Extended Data Fig. 6a), and down-regulation of *pfap2-i* in *Δpfap2-mrp* parasites at 16 h.p.i. (Fig. 3d). It has been reported that PfAP2-I binds the *pfap2-mrp* and its own promoter at 40 h.p.i.^30^, and in this study, we observed that PfAP2-MRP binds to its own promoter at both 16 h.p.i. and 40 h.p.i. (Extended Data Fig. 6b). Together, this information suggests that PfAP2-MRP autoregulates its expression at both 16 h.p.i. and 40 h.p.i., positively regulates the expression of *pfap2-i* at16 h.p.i.; PfAP2-I autoregulates its expression at 40 h.p.i. and *pfap2-mrp* expression might be additionally controlled by PfAP2-I at ∼ 40 h.p.i (combinatorial regulation; Extended Data Fig. 6c).

We found that PfAP2-MRP bound to the same promoter region of many genes as PfAP2-I and PfAP2-G, two proteins implicated in invasion^30^ and gametocytogenesis^31^, respectively (Extended Data Fig. 6d and 6e, Supplementary Data 3), suggesting a complex interplay of ApiAP2 DNA-binding proteins in control of gene expression (Supplementary discussion). Genes with promoters bound by PfAP2-MRP, PfAP2-I, and PfAP2-G include *gap45*; *msp1* and -*9*; *dblmsp*; e*tramp 4 -10.2* and *Pfap2-g* (Extended Data Fig. 6e).

### PfAP2-MRP is an indirect regulator of most invasion- and egress-associated genes

PfAP2-MRP binds to the promoters of only few invasion- and egress-associated genes at 40 h.p.i. (Extended Data Fig. 7 and Supplementary Data 3). These genes were significantly (FDR ≤ 0.05; log_2_ fold change ≤ -1) or weakly (log_2_ fold change between – 0.5 to -1) down-regulated in *Δpfap2-mrp* parasites at 40 h.p.i. compared to controls (Supplementary Data 1), suggesting that PfAP2-MRP directly regulates these genes. However, PfAP2-MRP was not bound to the promoters of most top down-regulated invasion- and egress-associated genes (Fig. 2b, Supplementary Data 1) except for *msa180*, zinc finger protein (PF3D7_0818100), *sub1* and *msp6* (Supplementary Data 3). These results suggest that the consequences of PfAP2-MRP knockout are both direct and indirect on gene expression.

### PfAP2-MRP binds to conserved sequence motifs

Analysis of the sequences bound by PfAP2-MRP at 16 h.p.i. and at 40 h.p.i. identified RCATGCR (6.1e^-36^, Fig. 4a); and GTGCR (8.7e^-28^, Fig. 4b) and RCATGCA (8.9e^-12^, Fig. 4c) as the most significantly enriched motifs respectively, excluding highly degenerate motifs. GTGCR is very similar to the motif bound by PfAP2-I *in vitro*^30^ (*P*= 4.7e^-03^, Fig. 4d), consistent with the observation that PfAP2-MRP binds many sequences bound by PfAP2-I (Extended Data Fig. 7d,e). RCATGCR (identified at 16 h.p.i.) is very similar to RCATGCA (identified at 40 h.p.i., *P*=6.97e-5, Fig. 4c) suggesting that PfAP2-MRP binding is sequence-specific with dynamic binding to similar sequence motifs in different genomic regions at different IDC stages (Fig. 4e-g and Supplementary Data 3). Such differential binding might be due to differences in PfAP2-MRP accessibility to different genomic regions, such as that conferred by differential chromatin structure^5^.

**Fig. 4.**
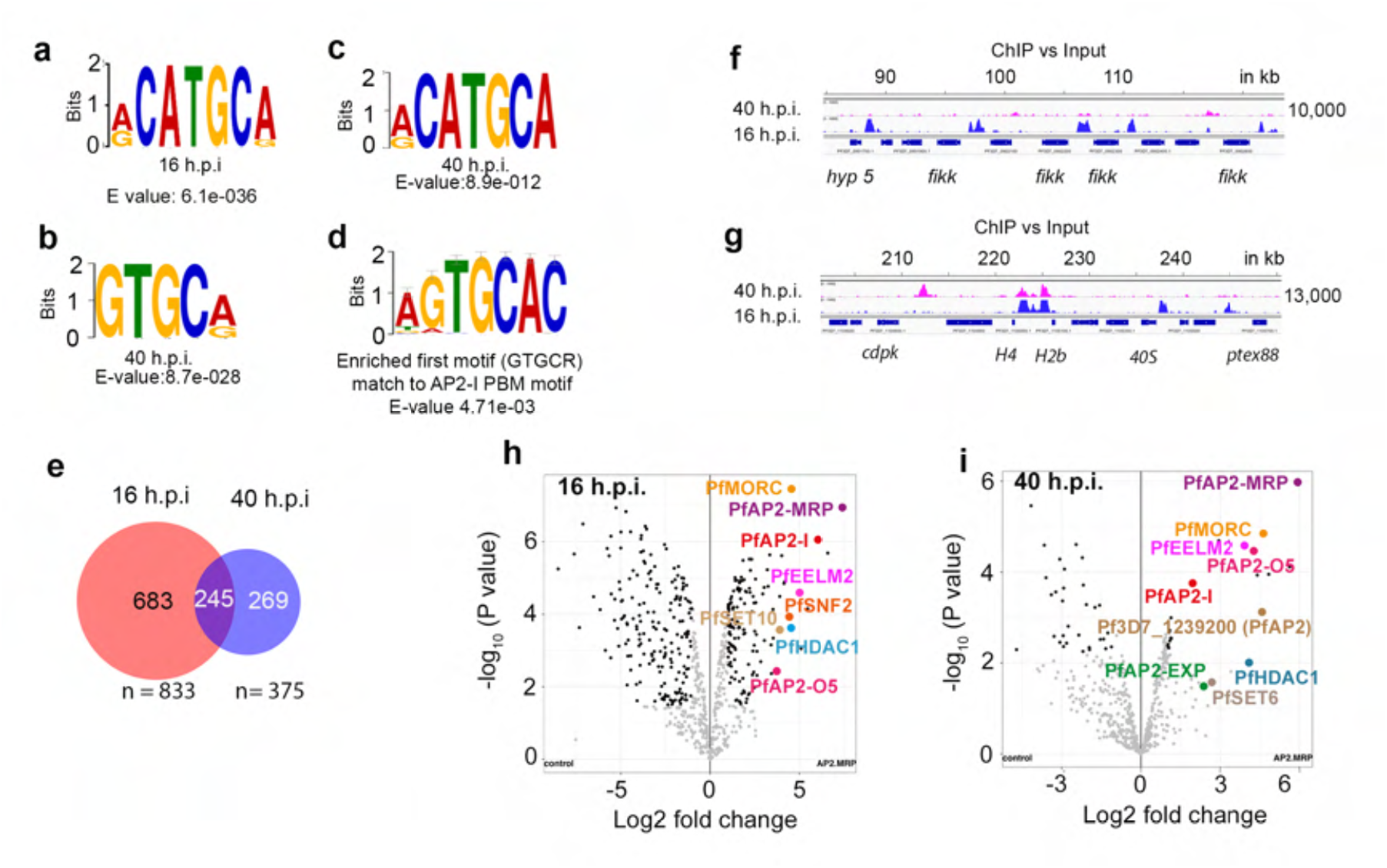
PfAP2-MRP binds to specific DNA motifs and associates with chromatin remodelers. **a,** Most significantly enriched motif bound by PfAP2-MRP at 16 h.p.i. **b-c**, The two most significantly enriched motifs bound by PfAP2-MRP at 40 h.p.i. **d**, The most enriched motif (panel **c**) at 40 h.p.i. is similar to the PfAP2-I binding motif **e**, The numbers of genes with promoters bound by PfAP2-MRP at either 16 h.p.i., 40 h.p.i. or both. **f**-**g**, Genomic regions uniquely bound by PfAP2-MRP at 16 h.p.i. (**f**) or at both 16 and 40 h.p.i. (**g**). X-axis shows the genomic position and numbers on the right, show the enrichment score. **h-i**, Label-free quantitative proteomic analysis of *P. falciparum* proteins enriched in PfAP2-MRP immunoprecipitates at 16h.p.i. and 40 h.p.i.

### PfAP2-MRP associates with known and putative epigenetic regulators

Since PfAP2-MRP activates or represses the expression of most gene families under epigenetic control and coding for clonally variant antigenic proteins, we proposed that PfAP2-MRP recruits epigenetic regulator(s) to modulate target genes. In support of this hypothesis, immunoprecipitation and mass spectrometry of PfAP2-MRP-3HA protein complexes identified several known or putative histone modifiers and chromatin remodelers, together with five other PfAP2 DNA binding proteins (Fig. 4h,i and Supplementary Data 5). While PfAP2MRP-associated histone modifiers such as PfSET10, PfSET6, and chromatin remodeler PfISWI proteins are transcriptional activators^32–34^, PfMORC-a putative chromatin remodeler along with PfHDAC1-a histone deacetylase are associated with repressed genes and are regarded as transcriptional repressors^35, 36^. Many of the proteins associated with PfAP2-MRP were present at both 16 and 40 h.p.i., but others were unique to one time point. The association of PfAP2-I, PfAP2-MRP and ISWI^30^ is consistent with the observed overlap between PfAP2-MRP and PfAP2-I binding sites and the significant similarity between the enriched motifs (Extended Data Fig. 7d,e).

### Depletion of PfAP2-MRP increases chromatin accessibility

While the relationship between chromatin organization and gene regulation remains unclear, parasite chromatin is organized into euchromatin and heterochromatin clusters and can act as a scaffold facilitating gene expression^37–39^. Furthermore, considering the strong effect of disruption of *pfap2-mrp* on genes associated with heterochromatin regions, we explored the effect of *pfap2-mrp* deletion on chromatin structure. We therefore performed comparative analysis of the chromatin conformation landscape using whole-genome chromosome conformation capture (Hi-C) followed by deep sequencing from either the tagged or the *Δpfap2-mrp* strains at 16 and 40 h.p.i. Correlation analysis of Hi-C replicates indicated a clear reproducibility at 16 and 40 h.p.i. where overall intra-chromosomal and interchromosomal interaction matrices appeared largely unchanged for each strain (Fig. 5a, and Extended Data Fig. 8). However, comparative analysis between the tagged and the *Δpfap2-mrp* lines revealed a slight reduction compared to background in long distance interactions and heterochromatin clusters in *Δpfap2-mrp* (Fig. 5b, and Extended Data Fig. 8) at 16 h.p.i. The effect was detected more globally at 40 h.p.i. Genome-wide mapping of interaction changes revealed a reduction in interaction frequency between telomere ends including internal *var* genes (Fig. 5c), consistent with a reduced clustering of the heterochromatin. Further support was provided by genome-wide 3D modeling on these two strains wherein we see clear separation of telomeric clustering in *Δpfap2-mrp* and overall expansion of the chromatin structure (Fig. 5d). We concluded that chromatin compaction is impaired in *pfap2-mrp* deficient parasites resulting in increased chromatin accessibility and up-regulation of *var* genes and genes involved in sexual differentiation. Such an effect could explain the partial disconnect between RNA-seq and ChIP-seq data observed in our study at both 16 and 40 h.p.i.

**Fig. 5.**
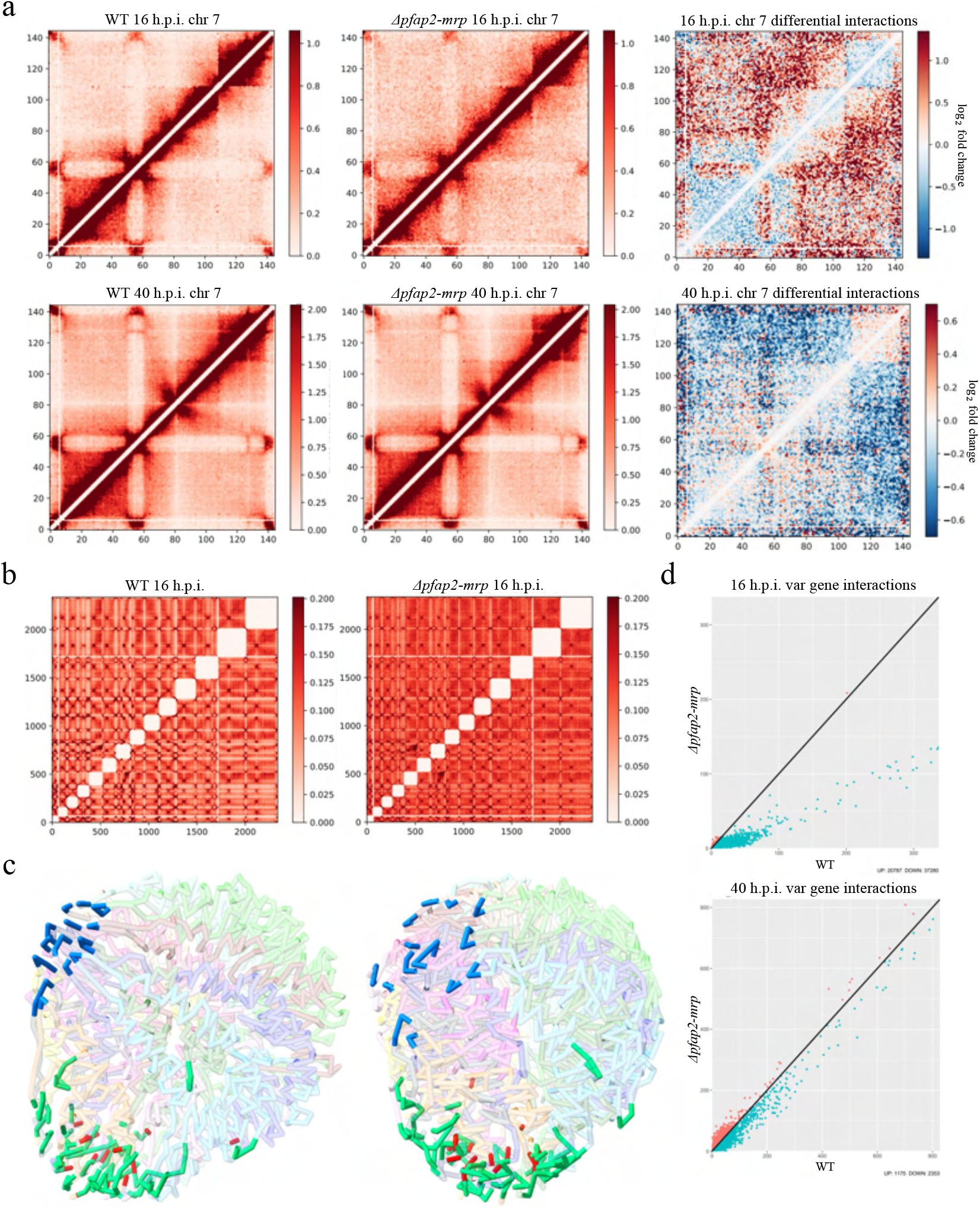
Depletion of PfAP2-MRP increases chromatin accessibility. **a,** Chromatin contact count heatmap of chromosome 7 at 16 h.p.i. (top) and 40 h.p.i (bottom) for the WT (left) and *Δpfap2-mrp* (middle), as well as the log_!_ fold change in interactions (right) between the WT and *Δpfap2-mrp*. Blue indicates a loss of interactions and red indicates an increase of interactions *Δpfap2-mrp* over WT. **b,** Whole-genome interchromosomal contact count heatmaps at 16 h.p.i. for the WT (left) and *Δpfap2-mrp* (right). Chromosomes are sorted from left to right and bottom to top. Intrachromosomal interactions are removed. **c,** Whole-genome 3D chromatin models at 16 h.p.i. for the WT (left) and *Δpfap2-mrp* (right). Centromeres (blue), telomeres (green), and *var* genes (red) are enhanced to display differences between the two samples. **d,** Number of interactions between *var* gene containing bins.

## DISCUSSION

We provide multiple lines of evidence that PfAP2-MRP is a master regulator that controls key processes of malaria development and pathogenesis during the IDC. Functional deletion of *PfAP2-MRP* at time points corresponding to its two peaks of expression enabled us to show that each peak has separate essential roles. The first peak at 16 h.p.i. is critical for parasite development beyond the trophozoite stage while the second peak is indispensable for merozoite development and egress (Extended Data Fig. 9). Through the use of RNA-seq, ChIP-seq and Hi-C, we show that PfAP2-MRP represses *vars* at both 40 h.p.i. and 16 h.p.i. by either interacting with their promoter or by regulating the chromosome accessibility that might contribute to the maintenance of chromatin compaction. scRNA-seq experiments further revealed the activation of most silenced *var* genes at 16 h.p.i. in Δ*PfAP2-MRP* parasites at a single nucleus level. The FACS-based experiment with pooled immune sera from a malaria-endemic region indicated greater immune recognition of the translated transcripts of multiple *var* genes as PfEMP1 proteins on the surface of iRBCs in the Δ*PfAP2-MRP* parasites. Taken together, these results suggest that PfAP2-MRP plays a crucial role in maintaining the mutually exclusive expression pattern of the *var* gene family by silencing most *var* genes. Regulation of *var* gene expression involves many chromatin-associated proteins^34^, and many of them were found to be associated with PfAP2-MRP, suggesting that it is a recruiter of these proteins (Supplementary discussion).

Disruption of PfAP2-MRP expression at 40 h.p.i. was sufficient to down-regulate genes associated with merozoite development, egress and invasion processes. Using mass spectrometry, we and other have demonstrated that PfAP2-MRP forms complexes with other Api-AP2s and chromatin-associated proteins, indicating that PfAP2-MRP-regulated genes may be under the combinatorial control of multiple regulatory factors (Supplementary Discussion). We showed that PfAP2-MRP and PfAP2-I^30^ bind to the same promoter regions of many invasion and egress-associated genes. Our combined results from RNA-seq and ChIP-seq experiments as well as IP-mass spectrometry indicate that a positive feedback loop-based transcriptional regulatory network exists between these two transcription factors. We observed over-expression of many early gametocyte marker genes in *Δpfap2-mrp* parasites in both 16 and 40 h.p.i. stages. This up-regulation of sexual genes may be explained by the decrease in heterochromatin clustering as well as a direct interaction with critical AP2 transcription factors. Overall, our result indicates that as an essential positive regulator of asexual growth, PfAP2-MRP has evolved to prevent sexual commitment by repressing early gametocyte marker genes critical for sexual conversion (Supplementary discussion).

In addition to enrichment of egress and invasion-associated genes, we observed many genes encoding hypothetical proteins of unknown function in the list of most down-regulated genes at 40 h.p.i. (n=50). Based on the principle of ‘guilt-by-association’^40^, we suggest that most of the top down-regulated genes encode hypothetical proteins that are previously unidentified components of the egress and invasion pathways. For example, we note that four of these genes encode hypothetical proteins (Pf3D7_1014100, Pf3D7_0210600, PF3D7_0308300, PF3D7_0507400) that have recently been shown to be important in invasion and egress pathways^41–44^ are also down regulated in our study. Future studies will be required to identify downstream regulator(s) of many other invasion and egress-associated genes. It is most likely that the functional characterization of PfAP2-MRP-associated proteins will elucidate their specific roles in chromatin remodeling, histone modification, and transcription.

In conclusion, this study establishes that several important processes in malaria development and pathogenicity are controlled by a master regulator, PfAP2-MRP. This paves the way for further understanding of the regulatory mechanisms controlling gene expression. It may also facilitate novel therapeutic strategies, such as the use of *ϕ..pfap2-mrp* parasites as a live anti-disease vaccine that expresses most of the PfEMP1 repertoire to elicit antibody responses that will decrease severe malaria-associated disease.

## METHODS

### Parasite culture, maintenance, synchronization, and transfection

The DiCre-expressing *P. falciparum* clone II3^13^ was maintained in human A+ erythrocytes at 37 °C in RPMI 1640 medium containing AlbumaxII (Invitrogen) supplemented with 2 mM L-glutamine. Parasites were either synchronized by sorbitol treatment or by purifying mature schizont stages using 70% (v/v) isotonic Percoll (GE Healthcare Life Sciences) before allowing reinvasion to occur, followed by sorbitol treatment. For transfection of plasmid constructs, purified Percoll-enriched mature schizonts (∼20 μl packed volume) were suspended in 100 μl of P3 primary cell solution (Lonza) containing 60 μg of linearized repair plasmid DNA (repair plasmid 1 and 2 separately in two separate transfection experiments) and 20 μg of pDC2 plasmid with the required cloned guide DNA and electroporated using an Amaxa 4D-Nucleofactor (Lonza), using program FP158 as previously described^45^. Drug selection was applied ∼20 h after transfection with 5 nM WR99210 (a kind gift of Jacobus Pharmaceuticals) for four days. Once sustained growth of drug-resistant transgenic parasites was observed, the cultures were treated with 1 μM 5-fluorocytosine (5-FC) provided as clinical Ancotil (MEDA) for four days. Transgenic parasite clones PfAP2-MRP:loxP and PfAP2-MRP-3HA:loxP were obtained by limiting dilution cloning in microplates. DiCre recombinase-mediated excision was induced by adding rapamycin to the culture at a final concentration of 10 nM.

### Strategy to perform conditional knockout of PfAP2-MRP

To excise exon 2 of *pfap2-mrp,* which contains the DNA-binding AP2 domain and nuclear localization signal, the endogenous exon 2 of *pfap2-mrp* in the II3 DiCre-expressing *P. falciparum* clone was replaced with transgenic, “floxed” and HA-tagged and non-tagged forms of the exon 2 using two sequential Cas9-mediated genome editing procedures. In the first step, the single intron of *pfap2-mrp* was replaced with *sera2*-loxPint^46^. The repair plasmid (repair plasmid 1) was synthesized commercially (GeneArt; Life Technologies) with recodonised sequences in the 2986–3024 bp and 3273–3305 bp regions of the *pfap2-mrp* and 400 bp homology arms flanking the 5’ and 3’ regions of the intron. In the second step, cloned parasites with the integrated *sera2-loxPint* were used to introduce a further loxP immediately after the stop codon of the *pfap2-mrp* gene. The repair plasmid for this (repair plasmid 2) contained recodonised sequences in the gene’s 6261–6330 bp region, a *XmaI* restriction enzyme site followed by a loxP sequence just after the TAA stop codon, and 400 bp homology regions.

A triple HA-tagged version of the PfAP2-MRP was also prepared. For this, the stop codon was removed from the donor sequence (repair plasmid 2) by PCR amplification using primer pairs (Oligos P1and P2) and digested with *NotI* and *XmaI* restriction enzymes. To remove the sequence with the stop codon from the plasmid backbone, repair plasmid 2 was also digested with *Not1* and *XmaI* restriction enzymes. Digested, amplified sequence without stop codon and digested plasmid backbone were ligated to get the repair plasmid 2 without stop codon after the PfAP2-MRP coding sequence. A triple HA tag sequence was PCR amplified from pFCSS plasmid with *XmaI* sites on both the sides of the HA tag sequence using oligos P5 and P6. *XmaI* digested repair plasmid 2 without stop codon was ligated with *XmaI* digested triple HA tag fragment to generate a triple HA-tagged encoding repair plasmid 2.

pDC2-Cas9-U6-hdhfr plasmid expressing spCas9 was used in this study. Guide RNA sequences were identified using Benchling. The TATTTATATTCTCAATTGAA and TTATATTCTCAATTGAATGG sequences targeting the 3’ end of the first exon, upstream of the TGG protospacer adjacent motif and 5’ end of the second exon upstream of the TGG protospacer, respectively, were cloned into the pDC2-Cas9-U6-hdhfr (pDC2 plasmid 1) and used to target Cas9 in the first Cas9-mediated editing step (**Fig. 1a**). The CCCTTCAATAGATTCGCACA sequence towards the 3’ end of the second exon, upstream of the CGG protospacer adjacent motif, was cloned into the pDC2-Cas9-U6-hdhfr (pDC2 plasmid 1) and used to target Cas9 in the second Cas9-mediated editing step. Sequences of all the oligos and primers used are listed in Supplementary Data 6.

### Flow cytometry-based quantification of parasitemia

Parasites were fixed in 0.1% glutaraldehyde, incubated for 30 minutes at 37 °C, then stored at 4 °C until further use. Glutaraldehyde fixed parasites were stained with SYBR green (diluted 1: 10,000) for 30 min at 37 °C, then parasitemia was determined using an BD LSR Fortessa flow cytometer (BD Biosciences, San Jose, CA, USA).

### Invasion and growth assays

For invasion assay, the parasites were treated with DMSO or rapamycin at ∼16 h.p.i. in cycle 0 and mature schizonts in the same cycle were isolated using Percoll as described earlier and added to the culture at 5% parasitemia. Merozoites were allowed to invade for 4 h under either static or vigorous shaking (250 rpm) conditions. Four hours after adding mature schizonts to the culture, the parasitemia was measured by flow cytometry. Three biological replicates per condition were used.

### Electron Microscopy

Compound 2 treated and RAPA treated (parasites with disrupted second peak of *pfap2-mrp* expression) highly synchronized parasites were allowed to grow until they reached the segmented schizont stage. Then the infected RBCs were fixed with 2.5% glutaraldehyde in cacodylate buffer (0.1 M, pH 7.4) for 48 h. A first osmication was performed using reduced osmium (1:1 mixture of 2% osmium tetroxide and 3% potassium ferrocyanide) for 1 hour. After a quick wash with distilled water, a second osmication was performed with 2% osmium tetroxide for 30 min. Samples were washed 3x 5min in water and placed in 1% Uranyl acetate for 12 hours at 4°C. Before dehydration, samples were washed 3x 15 min in water. After pre-embedding in 1% agar, samples were dehydrated in an ethanol series (40% to 100%) and embedded in epoxy resin. Thin sections (100 to 150 nm thick) were collected on copper grids and contrasted with lead citrate. Imaging was performed using a transmission electron microscope (TEM) operating at 300 kV (Titan Cryo Twin, Thermo Fisher Scientific). Images were recorded on a 4k x 4k CCD camera (Gatan Inc.).

### Nucleic acid extraction and polymerase chain reaction

For DNA isolation, cells were pelleted and treated with 0.15% saponin in PBS for 10 min on ice, then washed twice with PBS. DNA was extracted from parasite pellets using DNeasy blood and tissue kit (Qiagen) following the manufacturer’s instructions. For diagnostic PCR to check clones, GoTaq (Promega) DNA master mix was used; for amplification of fragments (3xHA tag) used in construct design, Phusion high fidelity DNA polymerase (NEB) was used, and for amplification of fragments longer than > 3 kb Platinum taq HiFi polymerase (Invitrogen) was used.

### PAGE, immunoblotting and immunofluorescence

Ring-stage parasites were treated with DMSO or rapamycin, and subsequently, mature schizonts (>45 h.p.i) were purified using 70% Percoll, then treated with 0.15% saponin in PBS and washed twice with PBS. Schizonts pellets were lysed by adding sample lysis buffer (1% NP40, 0.1% SDS, 150 mM NaCl), and 5 µg protein of each sample were separated under reducing conditions on Bis-Tris NuPAGE polyacrylamide gels and transferred to nitrocellulose membranes by electroblotting. Blots were blocked overnight in 5% milk powder (w/v) in phosphate-buffered saline (PBS) containing 0.2% Tween-20. To detect 3xHA tagged PfAP2-MRP, the rat anti-HA mAb 3F10 (Sigma) was used at a 1:1000 dilution, followed by horseradish conjugated secondary antibody (1:2500). For the proteins tested, relevant primary antibodies were used (see below), then secondary HRP conjugated antibodies specific for mouse, rabbit, or rat IgG (Biorad) were used at a dilution of 1:2500. The signal was developed using Immobilon Western Chemoluminescent HRP Substrate (Merck Millipore) and detected using Hyperfilm ECL film (GE Healthcare).

For immunofluorescence, thin parasite films were prepared and fixed in 4% paraformaldehyde in PBS for 20 min at room temperature. The cells were permeabilized with 0.1% (v/v) Triton X100 in PBS for 5 min, then blocked with 3% bovine serum albumin (BSA) in PBS overnight at 4 °C before being probed with relevant primary antibodies. Secondary Alexa Fluor 488- or 594-conjugated antibodies specific for mouse, rabbit, or rat IgG (Invitrogen) were used at a 1:5000 dilution. Slides were examined using a Nikon Ni microscope with a 100x Plan Apo NA 1.45 objective; images were captured with an Orca Flash 4 digital camera and prepared with Nikon NIS Elements and Adobe Photoshop software.

### Antibodies

The following antibodies and dilutions were used for Western blots in these studies. Rabbit anti-EBA175 (1:10,000), rat anti-MyoA (1:1,000), rat anti-BiP (1:1,000), rabbit anti-PTRAMP (1:4,000), rabbit anti-ARO (1:1,000), rabbit anti-AMA1 (1:10,000), and rabbit anti-SUB1 (1:1,000). Antibodies used in immunofluorescence were rabbit anti-GAP45 (1:1,000) and rabbit anti-MSP7 (1:1,000). Anti-EBA175 was obtained from MR4 (beiresources.org/MR4Home), anti-BiP was provided by Dr. Ellen Knuepfer, anti-SUB1 was a generous gift from Prof. Mike Blackman (Francis Crick Institute), and anti-AMA1 was a generous gift from Bart Faber and Clemens Kocken from the Primate Research centre in Rijswijk. All other antibodies were generated in the Holder laboratory and are now held and freely available at NIBSC-CFAR (please contact with any inquiries).

### RNA extraction and strand-specific RNA-seq library preparation

Parasite cultures (∼0.5-2 ml depending on the asexual developmental stage) were pelleted at 2400 rpm for 3 min, lysed by adding 1ml of Trizol (Sigma), then immediately stored at -80 °C until further use. Total RNA was isolated from Trizol lysed parasites following the manufacturer’s instructions (Life Technologies). Strand-specific mRNA libraries were prepared from total RNA using TruSeq Stranded mRNA Sample Prep Kit LS (Illumina) according to the manufacturer’s instructions. Briefly, for each sample, 100-300 ng of total RNA was used to prepare the libraries. PolyA+ mRNA was captured from total RNA using oligo-T attached to magnetic beads. First-strand synthesis was performed using random primers followed by second-strand synthesis where dUTP was incorporated in place of dTTP to achieve strand-specificity. Double-stranded cDNA ends were ligated with adaptors, and the libraries were amplified by PCR for 15 cycles before sequencing the libraries on Illumina HiSeq-4000 platform with paired-end 150 bp read chemistry according to manufacturer’s instructions (Illumina).

### RNA-seq data processing and analysis

The quality of the raw reads was assessed using FASTQC (http://www.bioinformatics.babraham.ac.uk/projects/fastqc). Low-quality reads and Illumina adaptor sequences from the read ends were removed using TrimmomaticR^47^. Processed reads were mapped to the *P. falciparum* 3D7 reference genome (release 40 in PlasmoDB-http://www.plasmoddb.org) using Hisat2^48^ (V 2.1.0) with parameter “—rna-strandness FR”. Counts per feature were estimated using FeatureCounts^49^. Raw read counts data were converted to counts per million (cpm), and genes were excluded if they failed to achieve a cpm value of 1 in at least one of the three replicates performed. Library sizes were scale-normalized by the TMM method using EdgeR software^50^ and further subjected to linear model analysis using the voom function in the limma package^51^. Differential expression analysis was performed using DeSeq2^52^. Genes with a false discovery rate corrected *P* value (Benjamini-Hochberg procedure) < 0.05 and log2 fold change ≥ 1 or ≤ -1 were considered as up-regulated or down regulated, respectively.

### Real-time quantitative PCR

Total RNA was treated with DNAase (TURBO DNase, Cat No. AM2238, Invitrogen) following the manufacturer’s instructions. The removal of DNA was confirmed by performing PCR using housekeeping genes. cDNA (1 µg) was prepared from total RNA using LunaScript cDNA synthesis mix (NEB), diluted 5 times before using it for qRT-PCR. mRNA expression levels were estimated on a Quant Studio 3 real-time qPCR machine (Applied Biosystems) using Fast SYBR green master mix (Applied Biosystems, Cat. No. 4385612). Seryl-tRNA ligase (PF3D7_0717700) was used as the internal control to normalize mRNA levels. Specific amplification of the PCR product was verified by dissociation curve analysis and relative quantities of mRNA calculated using the ΔΔCt Method^53^. PCR primers used in the real-time qRT-PCR experiment are listed in **Supplementary Data 6.** For the *var* gene qRT-PCR experiment, primer sets targeting individual *var* genes described elsewhere^54^ were used, including the primers targeting the seryl tRNA ligase gene (PF3D7_0717700) used as the internal control.

### PfAP2-MRP chromatin immunoprecipitation (AP2-MRP ChIP)

The ChIP assay was performed as described ^55^ with a few modifications. Parasite culture (50 ml) containing synchronized late-stage schizonts (∼ 40 h.p.i.) at ∼5% parasitemia was centrifuged at 900 g for 4 min, and the cells were washed once with PBS. To the cell pellet, 25 ml of 0.15% saponin in PBS was added and incubated on ice for 10 min, followed by washing twice with cold PBS. Parasites were crosslinked for 10 min by adding methanol-free formaldehyde at 1% final concentration and incubated for 10 min at 37 °C with occasional shaking. The crosslinking reaction was quenched by adding 1.25 M glycine to achieve a final concentration of 0.125 M and incubated at 37 °C for another 5 min. Parasites were centrifuged for 10 min at 3250 g at 4 °C, washed three times with DPBS, snap-frozen in liquid nitrogen, and stored at -80 °C until further use. Frozen formaldehyde-fixed parasites were thawed on ice for chromatin immunoprecipitation. One ml of nuclear extraction buffer (10 mM HEPES, 10 mM KCl, 0.1 mM EDTA, 0.1 mM EGTA, 1mM DTT, 1x EDTA-free protease inhibitor cocktail [Roche]) was added to the tubes containing thawed parasites and incubated on ice for 30 min. After the incubation, 10% NP-40 was added to reach a final concentration of 0.25%, and the parasites were lysed by passing through a 26^1/2^ G needle seven times. Parasite nuclei were collected by centrifuging at 2500*g* for 10 min at 4 °C. Shearing of chromatin was carried out using the Covaris Ultra Sonicator (E220) for 14 min with the following settings; 5% duty cycle, 140 intensity peak incident power, 200 cycles per burst to obtain fragment size of 200 to 600 bp. Insoluble materials were removed by centrifuging the sheared chromatin for 10 min at 13500 g at 4 °C. 30 µl of fragmented chromatin were stored as input at -80°C.

Fragmented chromatin was diluted 1:1 in ChIP dilution buffer (30 mM Tris-HCl pH 8.0, 0.1% SDS, 3 mM EDTA, 300 mM NaCl, 1% Triton X-100, EDTA-free protease inhibitor cocktail). Chromatin was incubated overnight with 6 µg of rabbit polyclonal anti-HA (Abcam no. ab9110) or, as control, the same amount of rabbit IgG isotype control (Cat. No. 10500c, Invitrogen). The antibody-protein complex was recovered with protein A coupled to magnetic beads (Dynabeads, Invitrogen, Cat No. 10002D), followed by extensive washes with low salt immune complex wash buffer, high salt immune complex wash buffer (washes done at 4 °C), and TE buffer (washes done at RT). Chromatin was eluted with elution buffer (1% SDS, 0.1 M NaHCO_3_) at 45 °C for 30 min with shaking. Immunoprecipitated chromatin and input were reverse crosslinked overnight at 45 °C by adding 5 M NaCl to a final concentration of 0.5 M. Samples were treated with RNaseA for 30 min at 37 °C followed by a 2 h incubation at 45 °C with proteinase K (final concentration 66 µg/ml). DNA was purified using ChIP DNA clean & concentrator (Zymo Research, Cat. No. D5205).

### PfAP2-MRP ChIP-sequencing and analysis

Libraries were prepared using NEBNext Ultra II DNA library kit following the manufacturer’s instructions until the step of adapter ligation (Adapters were diluted at 1: 20 ratio). Adapter ligated libraries were purified using AmpureXP beads. The libraries were amplified for a total of 6 PCR cycles (2 min at 98 °C initial denaturation; 6 cycles of 30 s at 98 °C, 50 s at 62 °C, final extension 5 min at 62 °C) using the KAPA HiFi HotStart Ready Mix (Kapa Biosystems). Amplified libraries were purified, and size selected for 350 bp inserts using AmpureXP beads and sequenced on the Illumina HiSeqX platform with 150 bp paired-end read layouts. Low-quality reads and Illumina adaptor sequences from the read ends were removed using Trimmomatic^47^. Quality trimmed reads were aligned to the *P. falciparum* genome (plasmodb.org, v3, release v32) using HiSat2. Duplicate reads were removed using samtools (markdup) ^56^. GC bias was corrected using deeptool’s correctGCBias tool^57^. For coverage plots of AP2-MRP 40 h.p.i. and 16 h.p.i. ChIP-seq experiments, deeptool’s bamCompare tool was used to normalize the read coverage per base of the genome position (option ‘-bs 1’) in the respective input and ChIP samples or IgG and ChIP samples to the total number of reads in each library (-- normalizeUsing RPKM). Normalized input coverage or IgG coverage per bin was subtracted from the ChIP values (option – operation subtract). Coverage plots were visualized using IGV genome browser^58^.

ChIP-Peaks (q-value cutoff < 0.05) were identified using macs2^59^ by comparing the input with ChIP or IgG with ChIP with default settings but without prior peak modeling (option ‘-nomodel’), the fragment size set to 200 bp (option ‘-extsize 200’) and the genome size (option ‘-g’) set to 233332839. Robust common peaks between replicates were identified using bedtools ‘intersect’ (option –f 0.30 –r)^60^. Peak annotation was carried out using Homer’s annotatePeaks.pl that assigned each peak with the nearest downstream gene. After intersecting, common peaks with peak score > 50 were kept for further analysis. Enrichment heatmaps and profile plots were generated using the deepTools computeMatrix and plotHeatmap tools.

### Processing of published PfAP2-I and PfAP2-G ChIP-seq data

PfAP2-I and PfAP2-G ChIP-seq published raw data^30, 31^ were downloaded from ENA and processed exactly as the ChIP-seq data for PfAP2-MRP. ChIP-Peaks (q-value cutoff < 0.05) were identified using macs2^59^ by comparing the input with ChIP for both the replicates of PfAP2-G and PfAP2-I. Robust common peaks between PfAP2-MRP and PfAP2-I; PfAP2-MRP and PfAP2-MRP and between all three were identified using bedtools ‘intersect’ (option –f 0.30 –r) to find peaks that overlapped at least 30%.

### Parasite sample preparation for single-cell RNA-seq

For 40 h.p.i. time-point, tightly synchronous parasites were enriched using 63% Percol, washed twice with incomplete RPMI medium, and processed immediately on the 10X Chromium controller (10X Genomics, Pleasanton, CA). For 16 h.p.i. time point parasites were stained with Mitotracker Deep Red FM (Life Technologies, #M22426) for FACS analysis and flow sorting, respectively. Briefly, 50 µl of SYBR Green I stained RBCs were analyzed on BD LSR Fortessa Flow Cytometer with High Throughput sampler (BD Biosciences, San Jose, CA, USA) using BD FACS Diva Software v6.2 and 488 laser excitation / 530 emission filter to determine the concentration of SYBR Green I positive cells per microliter. A BD Influx Cell Sorter (BD Biosciences, San Jose, CA, USA) with BD FACS Sortware v1.0.01 software was used to sort ∼40,000 MitoTracker Deep Red FM – positive RBC’s using a 70 µm nozzle, a 640 nm laser excitation / 670 nm emission filter and a pressure setting of 30 psi. Post-sorted cell concentration and quality were checked using a Countess® II Automated Cell Counter (Invitrogen) and FLoid™ Cell Imaging Station (ThermoFisher). Finally, labeled cells (i.e., SYBR Green I or MitoTracker Deep Red FM positive cells) were then loaded onto a 10X chip (Chip G) and processed immediately on the 10X Chromium controller (10X Genomics, Pleasanton, CA).

### Single-cell RNA-seq library preparation

Single-cell libraries were constructed using the 10X Genomics Chromium Next GEM Single Cell 3**ʹ** Reagent Kits v3.1 with Single Index Kit T Set A. Due to the extremely low RNA content of single-cell malaria parasite and the AT-rich genome (∼70% AT), modifications to the cDNA amplification and library preparation workflow were made accordingly. These modifications included – 30x cDNA amplification cycles; taking 50% cDNA as input into library generation; reducing fragmentation time to 2 minutes; and changing the extension time to 65 °C during index PCR. Individual library Qc was performed using the BioAnalyzer HS DNA Assay kit (Agilent).

### Sequencing of single-cell RNA-seq library

Library concentration was determined with the KAPA Library Quantification Kit (ROCHE) using the QuantStudio3 Real-Time PCR systems (ThermoFisher) and assessed for fragment size using the BioAnalyzer HS DNA Assay kit (Agilent). Following library pooling in equimolar concentrations, a total of 1.2 nM library was sequenced on the Illumina NovaSeq 6000 with SP flow cell using version 1.5 chemistry as follows: Read 1 – 28bp; Index read i7 – 8bp; Index read i5 – 0bp; and Read 2 – 91bp.

### Single-Cell Transcriptome alignment and read count estimation

The Droplet-based sequencing reads were aligned to the Hybrid genome of Human hg38 and *P. falciparum* pd37 (PlasmoDB-46_Pfalciparum3D7_Genome.fasta) to remove any human transcript contamination. This was achieved using CellRanger standard pipeline using --nosecondary flag. The raw gene count matrix was subjected to various single-cell preprocessing steps separately.

### Preprocessing and Normalization

Primarily, the reads mapping to human genes were removed, followed by the identification and removal of empty droplets using the emptyDrops() function from R package dropletUtils v1.12.3^61^. This function determines whether the RNA content of a cell barcode differs considerably from the ambient background RNA present in each sample. Cells with FDR <= 0.001 (Benjamini–Hochberg-corrected) were examined for subsequent investigation. The per-cell quality metrics were computed by the addPerCellQC function of the scuttle package v1.2.1^62^. The deconvolution approach in the computeSumFactors function of the Scran R package v1.20.1^63^ was utilized to normalize cell-specific biases. We kept the mitochondrial genes of *P falciparum* since the proportion of UMIs allocated to mitochondrial genes in both controls and knockouts were similar.

Further, the doublets identification was performed using the computeDoubletDensity function of the scDblFinder Bioconductor package v1.6.0 **[Germain P (2021). scDblFinder: scDblFinder. R package version 1.6.0,** https://github.com/plger/scDblFinder.]. This was achieved in three steps. a) The log normalization of counts was achieved using the logNormCounts function of scuttle package. b) The modelGeneVarByPoisson function of scran^63^ was then used to model the per-gene variance followed by c) doublet score calculation using the top 10% of highly variable genes (HVGs). We then cleaned our data based on a 95% quantile cut-off. Additionally, the standard functions of the Seurat package^64^ were also used to generate intuitive QC plots.

#### Cell type and infection stage recognition

The transcriptomic data at single-cell resolution about the malaria life cycle was obtained from the Malaria Cell Atlas. The phenotypic data for MCA was obtained from https://github.com/vhowick/MalariaCellAtlas/blob/master/Expression_Matrices/10X/pf1 0xIDC/pf10xIDC_pheno.csv^65^. We used the SingleR package v1.6.1 to transfer labels from the reference atlas to each cell in our data. It identifies marker genes for each stage in the reference atlas and uses them to compute assignment scores (based on the Spearman correlation across markers) for each cell in the test dataset against each label in the reference. The top 20 marker genes were identified using the Wilcoxon ranked sum test^66^

The time-series data of the intra-erythrocytic development cycle (IDC) was obtained from^14^. To transfer the time-series labels from Bulk RNASeq, we used a subset of samples for both 16h.p.i and 40h.p.i to prevent label misassignments. Each biological replicate was considered a single cell and markers were identified using the “classic” method of SingleR.

The correlation between cells of different stages was carried out using the CorrelateReference() function of the CHETAH R package v1.8.0^67^. Similarly, we performed an inter-sample correlation between the transcriptional profiles of samples from different time points.

### Integration of in-house data with MCA

The cleaned data was then integrated with Malaria Cell Atlas Single Cell data using standard scRNAseq integration workflow as described elsewhere^64^. Briefly, we create an “integrated” data assay but first identify the pairwise anchors using FindIntegrationAnchors followed by IntegrateData that exploit this anchor-set to combine the MCA and In-house data. Next, we ran the standard workflow on integrated assay including scaling, dimension reduction and clustering using parameters described in https://satijalab.org/seurat/articles/integration_introduction.html#integration-goals-1. The clusters were then manually annotated using the RenameIdents function of Seurat ^64^.

### *Var* gene expression per cell

A set of 61 *var* genes were used for their average expression calculation. The *var* gene expression was calculated using Raw counts (RNA assay data slot). *var* gene Expression calculation=sum of *var* gene expression/sum of Expression of all genes.

### PfAP2-MRP motif identification

Sequences from the commonly identified peaks from two replicates were extracted from the *P. falciparum* 3D7 genome using the bedtools ‘getfasta’. These retrieved sequences were then uploaded to the DREME server^68^ to identify significantly enriched motifs in the peak region. Tomtom^69^ was used to compare the de novo identified motifs to previously in silico discovered motifs^70^.

### Immunoprecipitation of PfAP2-MRP and identification of proteins by mass spectrometry

To identify the AP2 interacting proteins, the on beads digestion with trypsin approach was used after immunoprecipitation of AP2 complex. Before performing the on beads digestion, after TE buffer washes the immunoprecipitated complex was washed two times with exchange buffer (100 mM NaCl, Tris 50 mM pH 7.5) for 10 minutes at 4°C. Subsequently, the beads were resuspended in 100 μl of 100 mM Triethylammonium bicarbonate (TEAB), and reduction of bound proteins was done with 1mM DTT at 37°C for 30 minutes on thermo mixture with constant shaking at 750 rpm. The sample was brought to room temperature and alkylated in the dark with 3mM iodoacetamide for 45 minutes, the excess iodoacetamide was quenched with 3mM DDT for 10 minutes (Sigma Aldrich). Afterward, the sample was digested with 2.5μg trypsin (Promega) overnight on a thermo mixture with constant shaking at 1000 rpm. The resulting digested peptides were purified from beads, and trypsin digestion was stopped by adding TFA to a 2% final concentration. The acidified peptide was desalted using Sep-Pak C18 1 cc Vac Cartridge, 50 mg Sorbent per Cartridge (Waters). In brief, the Sep-Pak column was conditioned with 1 ml 100% methanol twice and equilibrated with 1 ml of 0.1% trifluoroacetic acid (TFA) twice. After that, the digested acidified peptides were loaded. The bound peptides were washed with 1 ml of 0.1% TFA twice and eluted with 300 μl of elution buffer (Acetonitrile 75% with 0.1% TFA in water) twice. The eluted peptides were dried in SpeedVac and kept at -80°C till further use.

### LC-MS analysis of peptides

The LC-MS analysis was performed on Q-Exactive HF mass spectrometer coupled with an UltiMate™ 3000 UHPLC (Thermo Fisher Scientific). The peptides were dissolvedin 0.1% formic acid (FA; Sigma Aldrich), and approximately 1μg of peptides was separated on an Acclaim PepMap^TM^ C18 column (75 μm I.D. X 250 mm, 2 μm particle sizes, 100 Å pore sizes) with a gradient of 5-35% mobile phase A and B respectively for 55 minutes, ramping up to 90% phase B for 5 minutes, 90% phase B for 5 minutes and the column was conditioned to 2% phase B for 10 minutes with the flow rate of 300 nl/min^-1^ (Phase A 0.1% FA, Phase B 99.9 % ACN with 0.1%FA). The peptides were introduced into the mass spectrometer through Nanospray Flex with an electrospray potential of 2.5 kV. Data were acquired in the Orbitrap at the resolution of 60,000 in the mass range of 350-16000 m/z with a maximum ion accumulation time set to 50 ms. The 20 most intense ions with a threshold of more than 1x e^6^ and having the multiple charges were further fragmented by using higher-energy collision dissociation (HCD) at 15000 resolution. The Dynamic exclusion for HCD fragmentation was 30 seconds. The maximum time for fragmented ion accumulation was 30 ms, a target value of 2.50xe^3^, the normalized collision energy at 28%, and the isolation width of 1.6. During the acquisition, the ion transfer tube temperature was set at 160°C, data was acquired in data-dependent acquisition (DDA) mode, and the total run time was 75 minutes.

### Identification, quantification, and statistical analysis of LC-MS data

Raw LC-MS data files from Q-Exactive HF were converted to .mgf files using Proteo Wizard MS covertgui 64 bit and analyzed using Mascot v2.4. The annotated protein sequence for *Plasmodium falciparum* was downloaded from https://plasmodb.org (Release 51). Trypsin was set as the enzyme of choice with maximum missed cleavage 1, fixed modification carbamidomethyl (K), variable modification deamidated and oxidation of N, Q, and M, respectively, with the peptide and fragment mass tolerance at 0.6 Da.

### ChIP qPCR

After de-crosslinking, ChIP DNA was purified using the Zymo ChIP DNA kit and quantified by Qubit HS DNA assay. The purified ChIP DNA was first diluted 20-fold in elution buffer and then analyzed by qPCR using the CFX-96 Biorad system. All ChIP primers used (Supplementary Data 6) were first checked using genomic DNA to determine specificity (based on a single peak in the melting curve) and efficiency. ChIP qPCR data were analyzed using the ΔΔCt method. *Pfap2-mrp* ChIP-qPCR results are expressed as a percentage of input. Three biological replicates of both samples and negative control (mock IP using IgG) were used for ChIP-qPCR experiments.

### Flow cytometry using pooled human serum

iRBCs with trophozoite stage parasites from cycle 1, treated with DMSO or rapamycin in cyle 0 to disrupt the first peak of *PfAP2-MRP* expression, were washed thrice with PBS supplemented with 0.1% BSA. iRBCS were either untreated or treated with Hiserum. When untreated, the same volume of 0.1% BSA in PBS was added and incubated for 30 mins at room temperature. Cells were washed thrice with 0.1% BSA in PBS, and all the samples were treated with sybr green (1x) and mouse anti-human IgG conjugated with Alexflour 647 (1:100 dilution, from BioLegend) for 30 mins at room temperature. After incubation, samples were washed thrice again with 0.1% BSA in PBS and analyzed on an BD LSR Fortessa flow cytometer (BD). Data were analyzed using FlowJo v9 software.

### In situ Hi-C

Parasite were crosslinked using 1.25% formaldehyde in warm 1X PBS for 25 min at 37 °C with rotation. Glycine was then added to a final concentration of 150 mM to quench the formaldehyde and incubated for 15 min at 37 °C and 15 min at 4 °C, both with rotation. Following centrifugation at 660 x g for 15 min at 4 °C, the pellet was resuspended in 5 volumes of ice-cold 1X PBS and incubated for 10 min at 4°C with rotation. After another centrifugation at 660 x g for 15 min at 4 °C the pellet was resuspended in 20 ml ice-cold 1X PBS. Several more washes in cold 1X PBS were used to clear cellular debris before resuspending in 1 ml 1X PBS and separated into multiple 1.5 ml tubes at a concentration of 1 x 10^8^ parasites per tube. The tubes were flash frozen in liquid nitrogen and stored at 80 °C before continuing with the rest of the in-situ Hi-C protocol^71^ using MboI restriction enzyme, with modifications to the standard protocol^72^. The final Hi-C libraries were sequenced using the Illumina NovaSeq 6000 using the S4 300 cycle flow cell for paired-end read libraries.

### Hi-C data processing and analysis

Paired-end HiC library reads were processed using HiC-Pro (Servant et al., 2015) with default parameters and mapping at 10kb resolution to the *P. falciparum* genome (release-50, plasmodb.org). The ICED-normalized interaction matrices output by HiC-Pro were interaction counts-per-million normalized before generating interaction heatmaps. All intra-bin contacts and contacts within a two-bin distance were set to 0 to enhance visualization and the color map per chromosome was scaled based on the minimum number of interactions in the highest 10% of interacting bins to aid in comparison between samples. Interaction matrices for the replicates were merged and differential interactions were identified by calculating the log_2_ fold change between the wild type and AP2-KO at each time point. Coordinate matrices generated by PASTIS^73^ were visualized as 3D chromatin models using ChimeraX^74^.

## Acknowledgements

The project was supported by a faculty baseline fund (BAS/1/1020-01-01) and a Competitive Research Grant (CRG) award from OSR (OSR-2018-CRG6-3392) from the King Abdullah University of Science and Technology (KAUST) to AP. A.A.H. is supported by the Francis Crick Institute (FC010097), which receives its core funding from Cancer Research UK (FC010097), the UK Medical Research Council (FC10097), and the Wellcome Trust (FC010097). This research was funded in part, by the Wellcome Trust [FC010097] and for Open Access, the author has applied a CC BY public copyright licence to any Author Accepted Manuscript version arising from this submission. It was also supported by the National Institutes of Allergy and Infectious Diseases and the National Institutes of Health (grant R01 AI136511 and R21 AI142506-01 to KLR) and the University of California, Riverside (NIFA-Hatch-225935 to KLR). The authors thank the staff of the Bioscience Core Laboratory in KAUST for sequencing bulk and single-cell RNA-seq libraries and FACS assays and all the members of the Holder lab at the Francis Crick Institute, London and Pathogen Genomics Lab at KAUST for assistance during the experiments. We also acknowledge Mohammed Almohammadi, Farzal Anwar from the blood transfusion centre associated with King Abdulaziz Medical City, Jeddah, Saudi Arabia for supplying us fresh RBCs and Sharif Hala from Infectious Disease Research Department, King Abdullah International Medical Research Center, Jeddah, Saudi Arabia for coordinating the logistics for the supply of RBCs to support part of the malaria culture work. HiSerum used in the FACS based experiment was a kind gift from Alister Craig at the Liverpool School of Tropical Medicine (LSTM).

## Data availability

The data sets generated in this study are available in the following databases:

- RNA-seq data: NCBI BioProject accession # GSE190342
- ScRNA-seq: NCBI BioProject accession # GSE191025
- ChIP-seq data: NCBI BioProject accession # GSE 190497
- Proteomics data: Pride accession number # PXD030308.
- Hi-C data: ENA BioProject accession number #PRJNA847684

The bulk RNA-seq, ScRNA-seq and ChIP-Seq datasets have been added under the super series GSE190519

## Author contributions

Conceptualization, A.K.S., and A.P.; Methodology, A.K.S., J.L.G., A.A.H., and A.P.; Investigation, A.K.S., J.L.G., M.G., R.P.S., M.S., I.I., T.M., R.N., Z.S., S.M., L.E., R.S., Y.O., R.S., F.B.R., S.A., A.D., A.F.K., J.M., W.F., E.K., I.G., D.J.P.F.; Analysis, A.K.S., J.L.G., R.S., S.A., T.L., I.G.; Writing the original draft, A.K.S.; Writing-Review and Editing, A.K.S., J.L.G., K.G.L.R., A.A.H and A.P.; Resources, A.A.H., and A.P.; Funding acquisition, A.A.H., and A.P.; Supervision, A.P. All authors read and approved the manuscript.

## Figures

**Extended Data Fig. 1.**
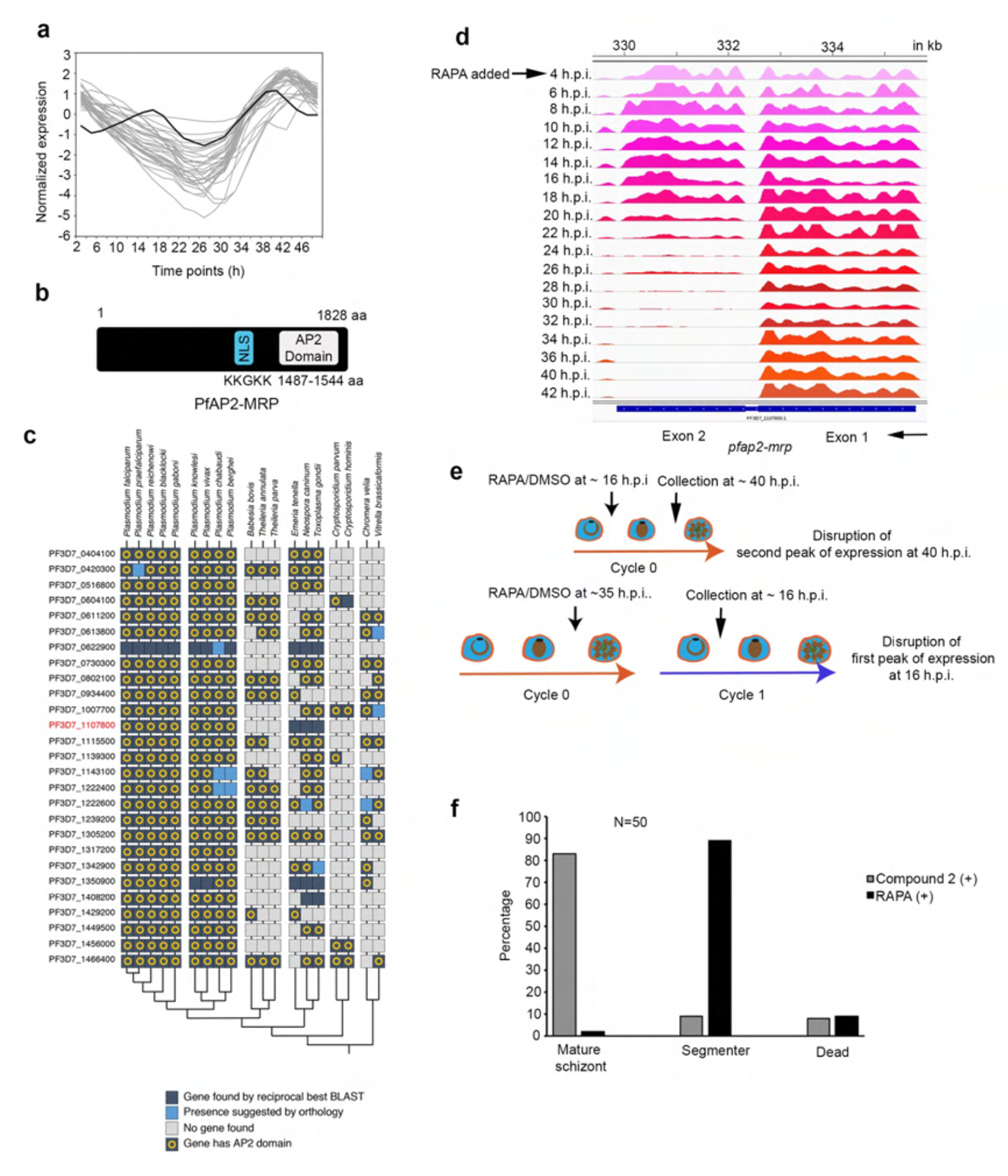
*pfap2-mrp* knockout affects merozoite development. **a,** Expression of *pfap2-mrp* (thick black line) and the top 40 down-regulated genes (grey lines) in *P. falciparum* line II3 over a 48-hour IDC^14^. **b,** The primary structure of PfAP2-MRP, with a nuclear localization signal (NLS) and a single AP2 DNA binding domain. Both NLS and DNA binding domain are encoded by exon 2 of the gene. **c**, A phylogenetic distribution of AP2 genes in the genus *Plasmodium* and related alveolates. **d**, RNA-seq data from different IDC time-points mapped to the *pfap2-mrp* locus; there is a drastic reduction in RNA-seq reads mapping to the second exon 16 hours after the addition of rapamycin i.e. at 20 h.p.i. **e**, Schematic showing rapamycin treatment schedule to disrupt either first or second peak of *pfap2-mrp* expression. **f**, The different developmental stages of the Compound 2- and RAPA-treated parasites (at 49 h.p.i.); 50 randomly selected iRBCs from each group were inspected and the parasites were categorized as either mature schizont, segmenter or dead.

**Extended Data Fig. 2.**
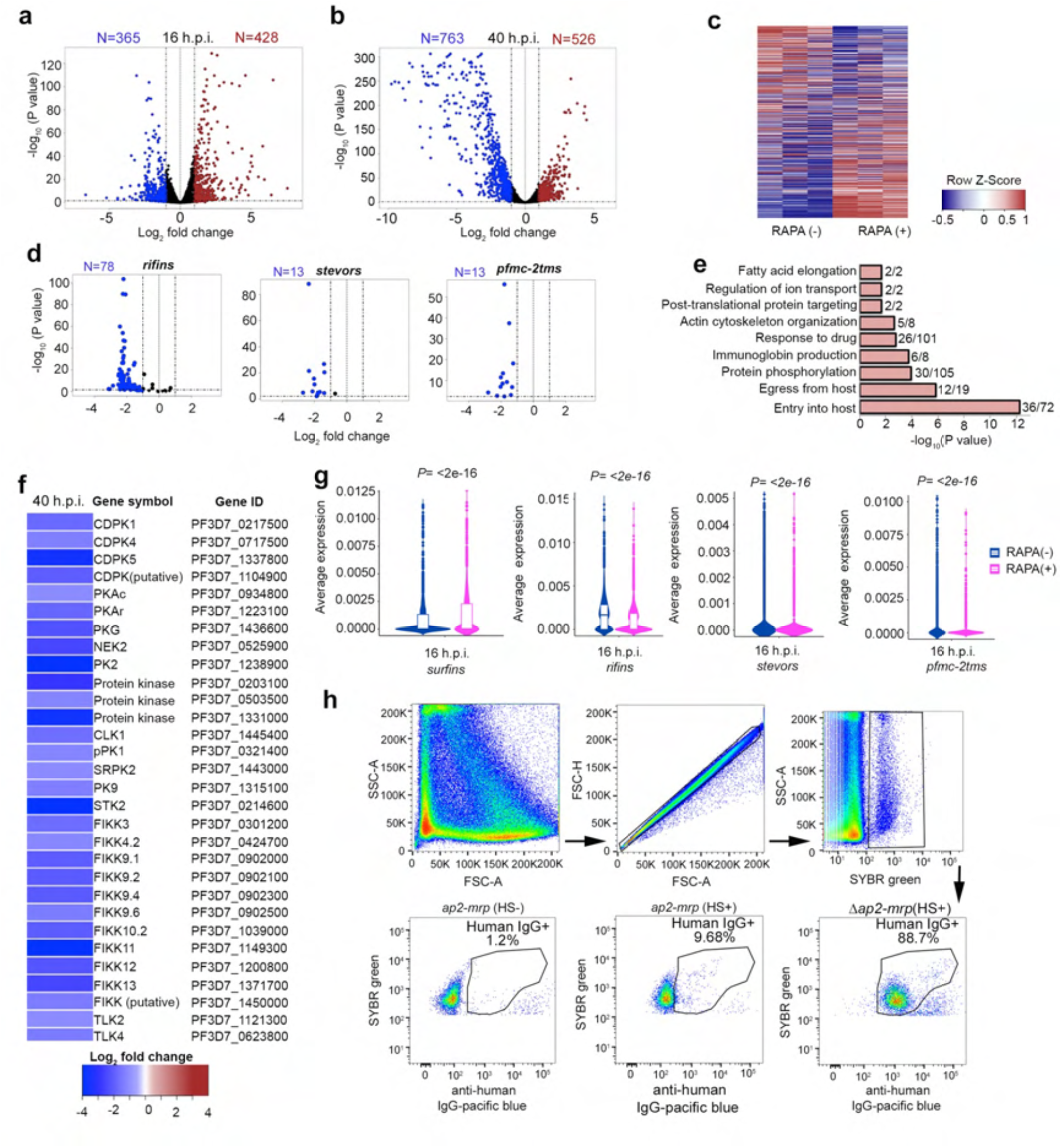
Disruption of *pfap2-mrp* deregulates malaria pathogenesis-associated genes. **a-b**, Volcano plots showing significantly differentially expressed genes in rapamycin-treated compared to control parasites at 16 h.p.i. (**a**) and 40 h.p.i. (**b**). **c**, Non-differentially expressed genes in treated [RAPA (+)] compared to control [RAPA (-)] parasites at 40 h.p.i., which had been reported to express at least 4-fold higher in schizont-stage parasites (> 35 h.p.i.) compared to early-stage parasites (< 35 h.p.i.)^5^. **d**, Expression of members of the gene families: *rifin*, *stevor* and *Pfmc-2tm* at 16 h.p.i.that encode antigenically variant proteins. **e**, Gene-ontology (GO) enrichment analysis of all genes down-regulated in rapamycin-treated compared with control parasites at 40 h.p.i. Shown is the number of genes down-regulated in treated parasites out of the total number of genes assigned to that specific GO term. **f**, Heatmap of the expression of all known and putative *P. falciparum* kinases downregulated following rapamycin treatment compared to controls, at 40 h.p.i. **g**, Violin plots of all *surfin*, *rifin, stevor* and *pfmc-2tm* expression per cell in treated [RAPA (+)] and control [RAPA (-)] parasites at 16 h.p.i.. **h**, FACS gating strategy for surface PfEMP1 expression detected by IgG binding from pooled serum of malaria-infected individuals. Cells were incubated either without (HS-) or with (HS+) pooled serum from malaria-infected individuals. The percentage of Human IgG+ iRBCs is the mean of three biological replicates. A total of 7,500 Sybr green positive events (iRBCs) per sample were analyzed.

**Extended Data Fig. 3.**
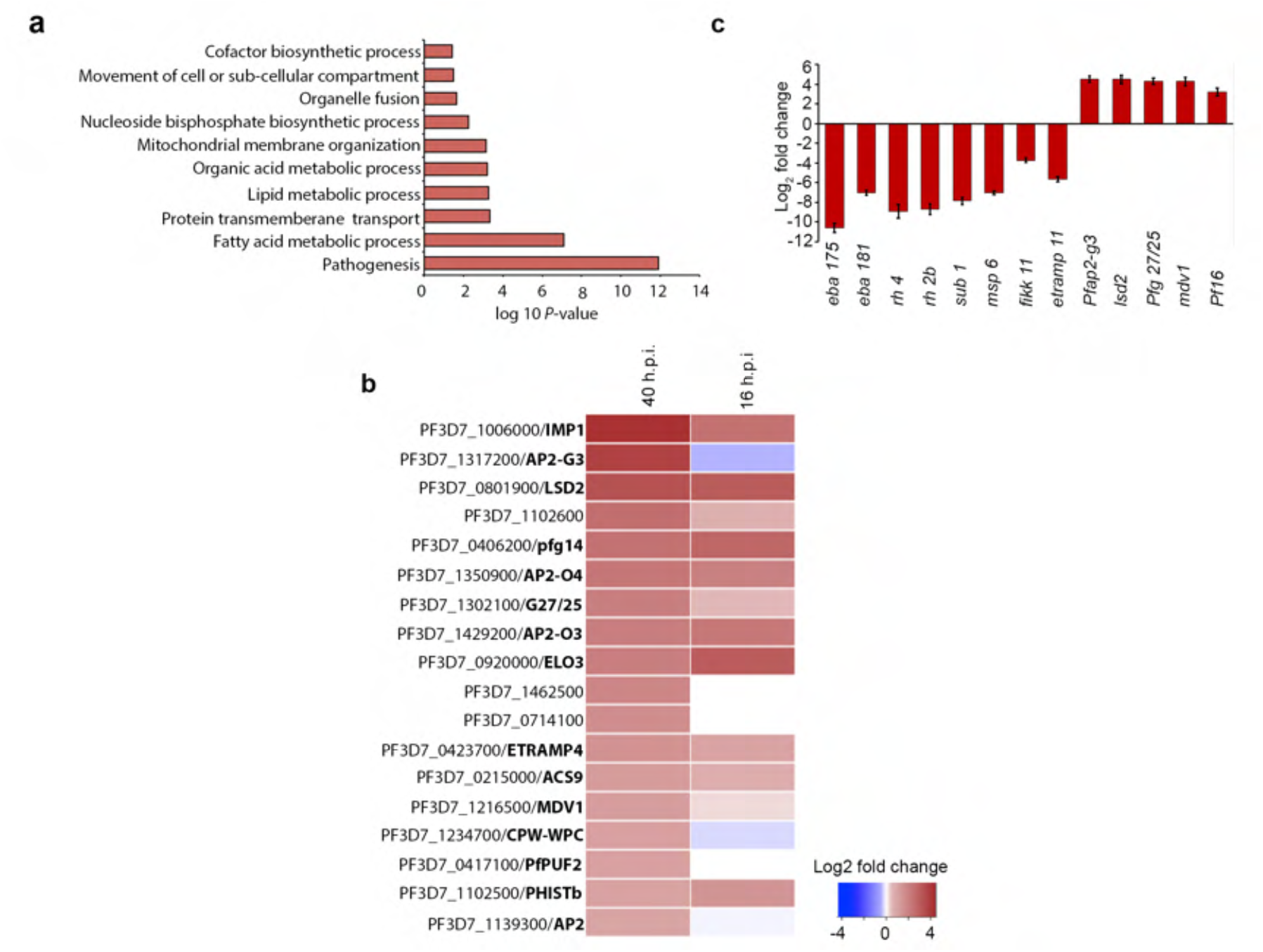
PfAP2-MRP is a repressor of early gametocyte marker genes. **a**, Gene ontology enrichment analysis of up-regulated genes in *Δpfap2-mrp* parasites at 40 h.p.i. **b**, Differential expression in *Δpfap2-mrp* parasites at 40 h.p.i. and 16 h.p.i. of genes that are known or putative early gametocyte markers. **c**, Differential expression of selected genes in RAPA-treated and control parasites measured using qRT-PCR, to validate RNA-seq data.

**Extended Data Fig. 4.**
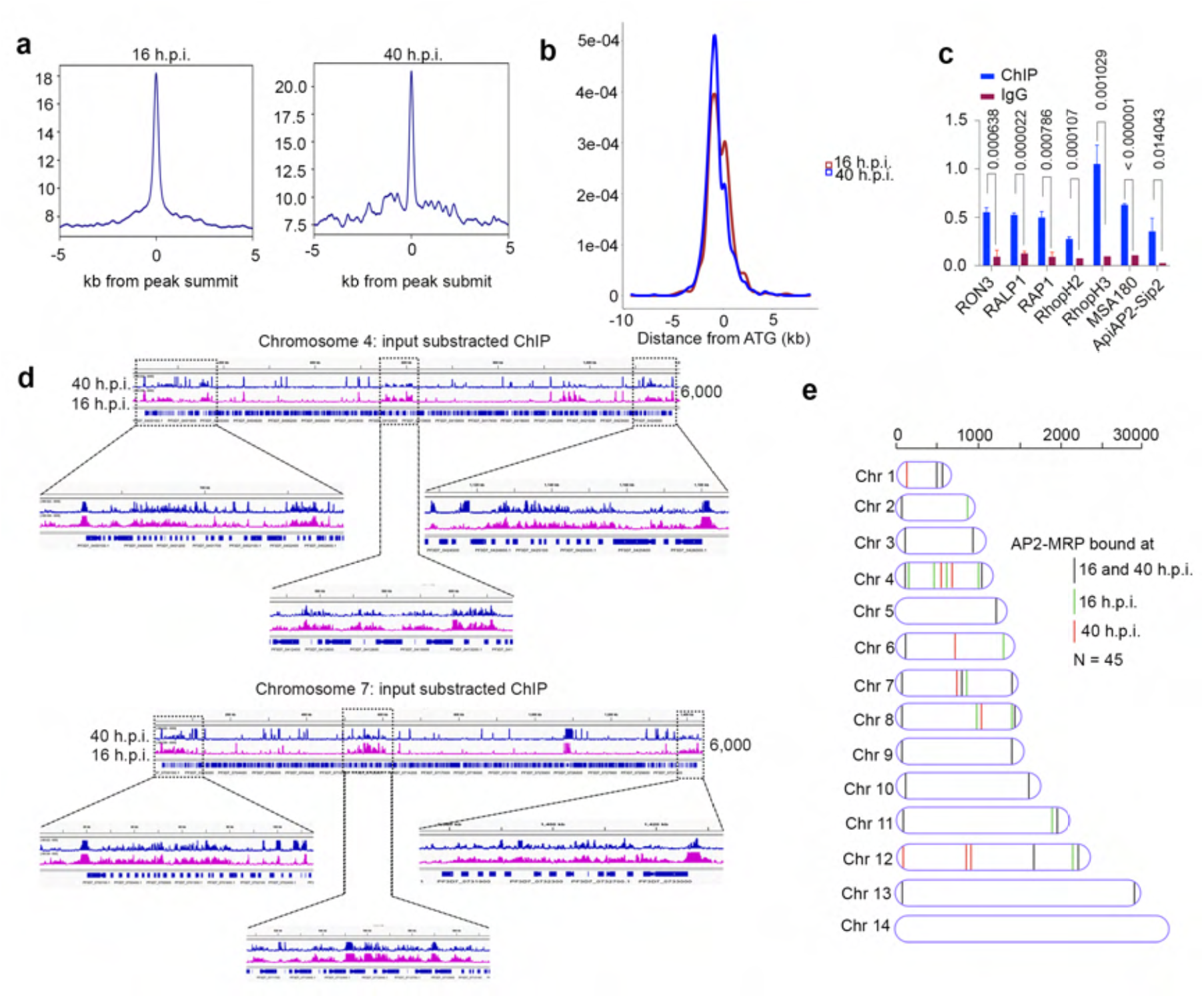
PfAP2-MRP binds to the putative promoter regions of *var* genes. **a**, Enrichment of PfAP2-MRP bound reads (from replicate 1) around the ± 5 kb region of peak summits, from 16 h.p.i. (left panel) and 40 h.p.i. (right panel) parasite samples. **b**, The position of ChIP-seq peak summits (common between two biological replicates) relative to the predicted ATG translational start codon, in 16 h.p.i. (red) and 40 h.p.i. (blue) parasites. **c**, Three independent PfAP2-MRP ChIP experiments followed by qPCR, were performed to validate ChIP-seq data, using selected PfAP2-MRP-bound promoter regions of genes from samples at 40 h.p.i. The bar-plot shows percent input (% Input) enrichment of PfAP2-MRP on target genes (mean ± SD of three independent experiments). IgG was used as the mock-treated control. *P*-values were calculated using a two-tailed t-test. **d**, Input subtracted ChIP peaks of PfAP2-MRP in chromosome 4 and 7 as representatives in both 16 and 40 h.p.i. stages. Also, zoomed in PfAP2-MRP bound central chromosomal and sub-telomeric heterochromatin regions are shown. X-axis shows the genomic position and numbers on the right show the enrichment score. **e**, Schematic diagram of the chromosomal position of *var* genes with promoters bound by PfAP2-MRP at either 16 h.p.i. (green), 40 h.p.i. (red) or at both stages (black).

**Extended Data Fig. 5.**
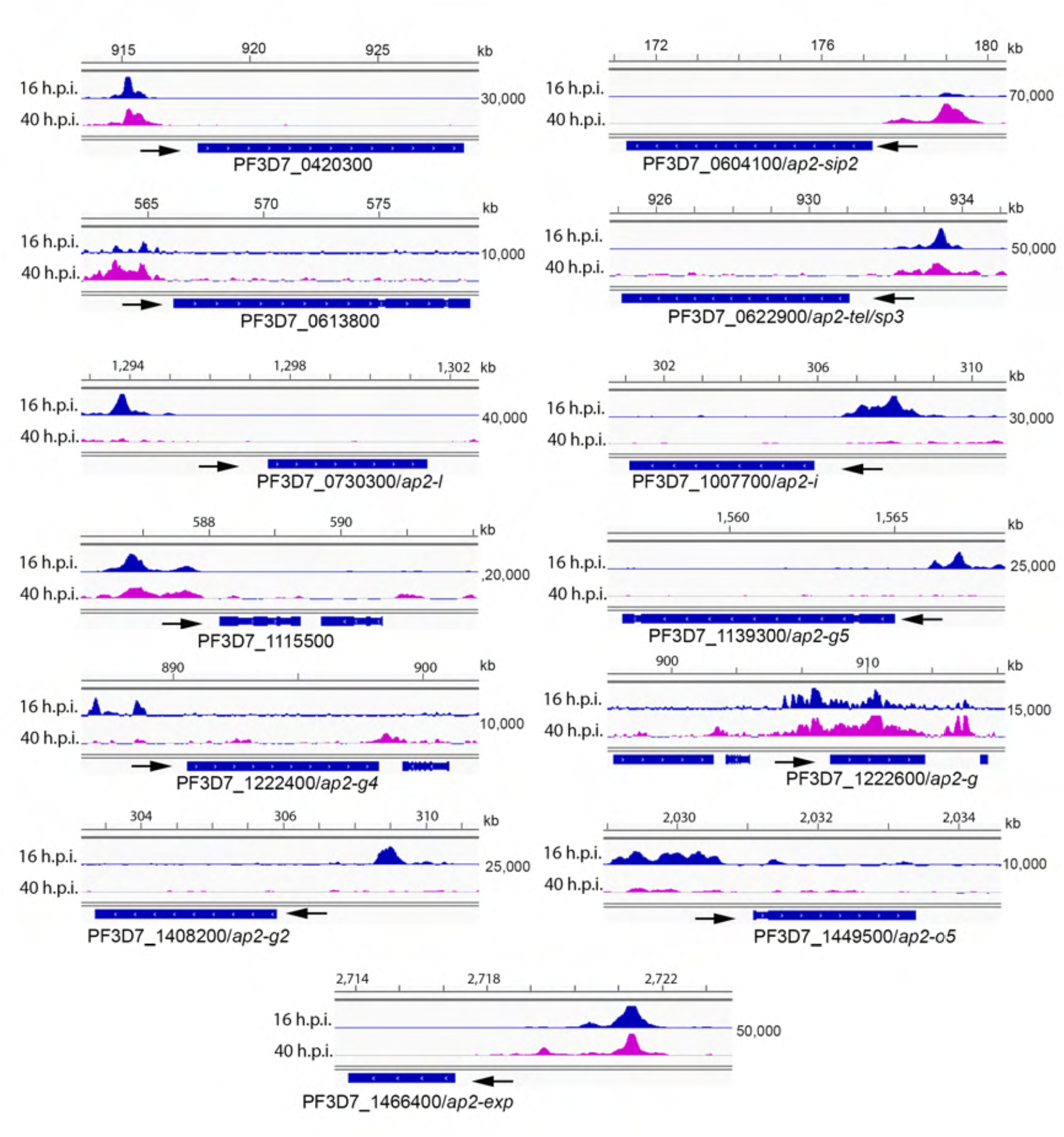
PfAP2-MRP binds to the putative promoter regions of multiple *apiap2* genes. Occupancy of PfAP2-MRP in the promoter region of *apiap2* genes. ChIP tracks show input subtracted PfAP2-MRP-ChIP from 16 and 40 h.p.i. parasites. Arrow marks show the direction of gene transcription X-axis shows the genomic position, and numbers on the right show the enrichment score.

**Extended Data Fig. 6.**
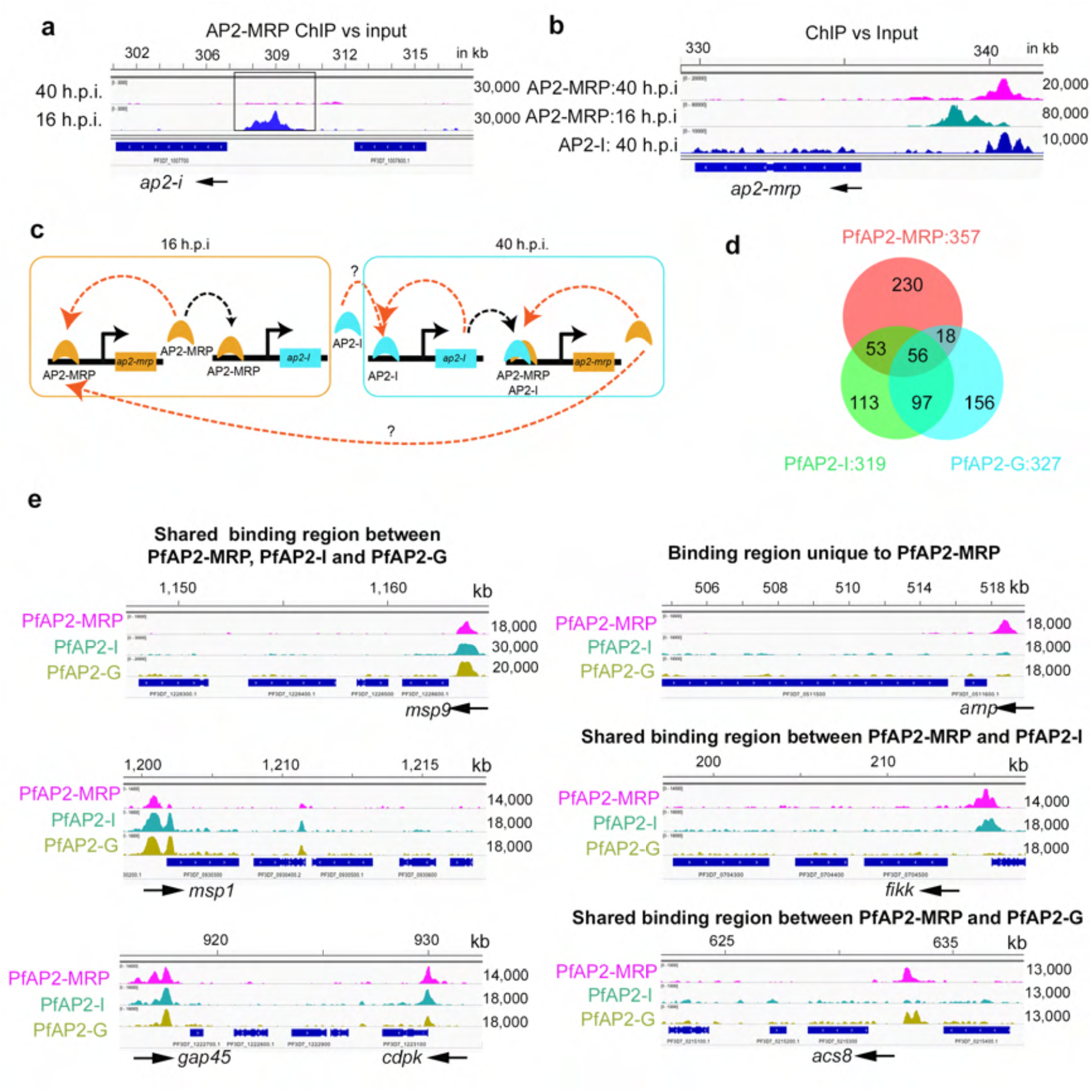
PfAP2-MRP, PfAP2-I and PfAP2-G binds to many common genomic regions. **a**, Occupancy of PfAP2-MRP in the promoter region of *pfap2-I* at 16 and 40 h.p.i. **b**, Occupancy of PfAP2-MRP and PfAP2-I in the promoter region of *pfap2-mrp* at 16 and 40 h.p.i. **c**, Schematic showing probable gene regulatory network between PfAP2-MRP and PfAP2-I. Orange arrows indicate the binding of protein to its gene promoter. **d**, Comparison of genes with promoters bound by PfAP2-MRP (red), PfAP2-I (green) and PfAP2-G (light blue) at 40 h.p.i. **e**, Input-subtracted ChIP-seq read coverage for exemplar genes with promoters that are either bound by all three AP2 proteins (left panel), uniquely by PfAP2-MRP, by both PfAP2-MRP and PfAP2-I, or by both PfAP2-MRP and PfAP2-G (right panel). Arrows show the direction of gene transcription. X-axes show the chromosomal position, and the numbers on the side show the peak enrichment score. X-axis shows the genomic position and numbers on the right show the enrichment score.

**Extended Data Fig. 7.**
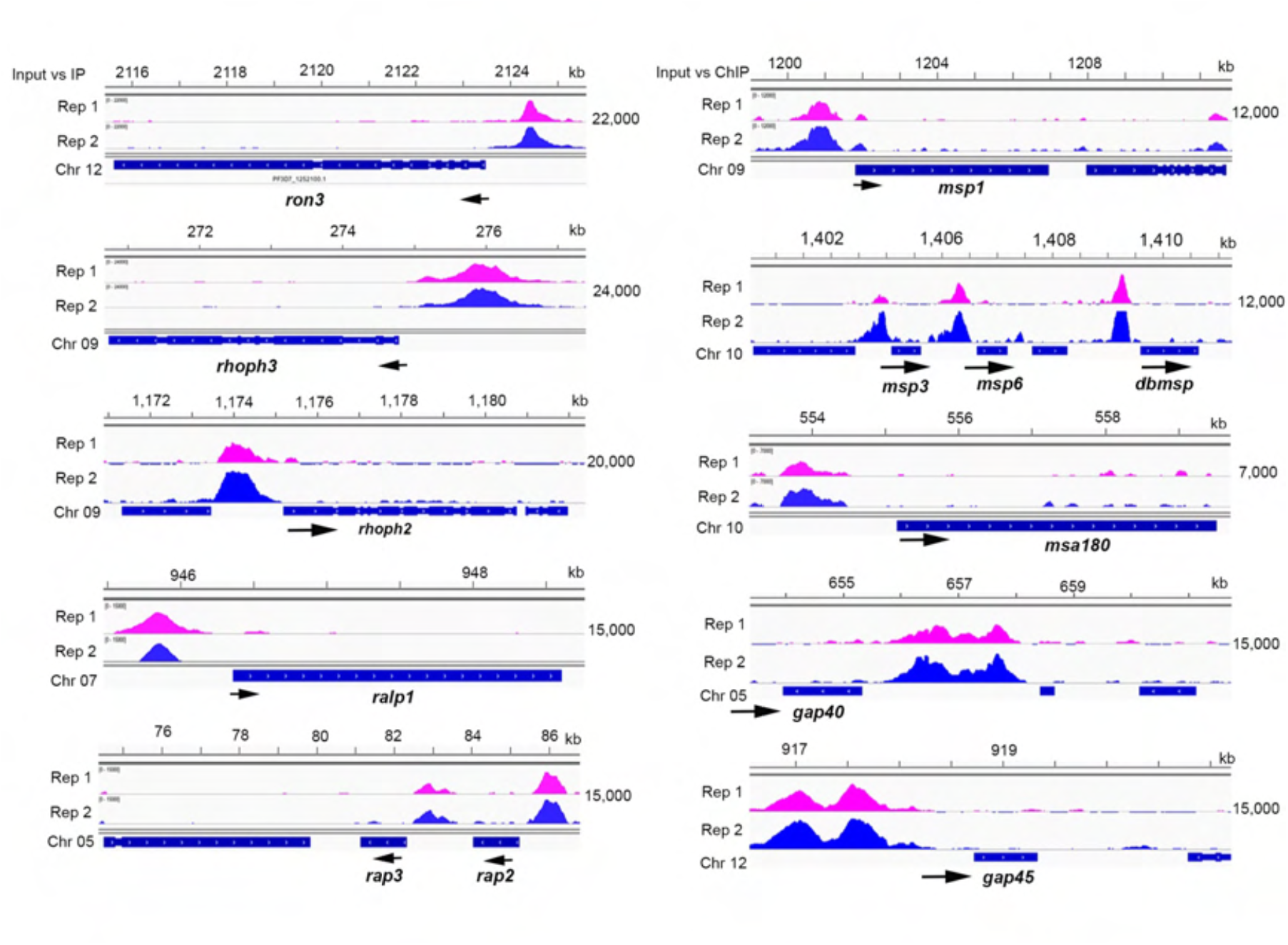
PfAP2-MRP binds to the putative promoter region of invasion-associated genes. Occupancy of PfAP2-MRP in the promoter region of different invasion associated genes. The ChIP tracks show two replicates with input subtracted from the PfAP2-MRP-ChIP data. Arrows indicate direction of transcription. X-axis shows the chromosomal position, and numbers on the right, show the enrichment score.

**Extended Data Fig. 8.**
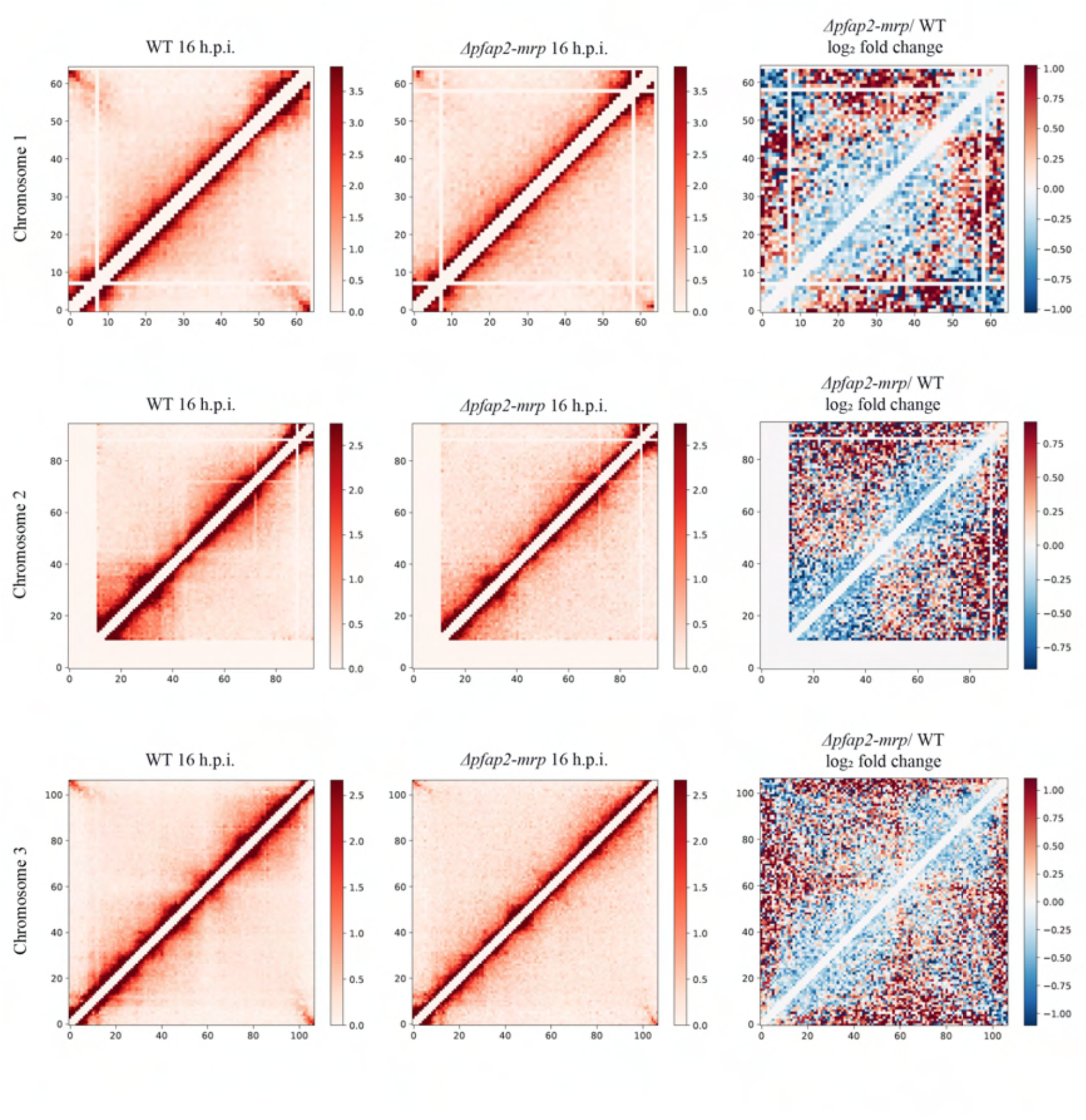
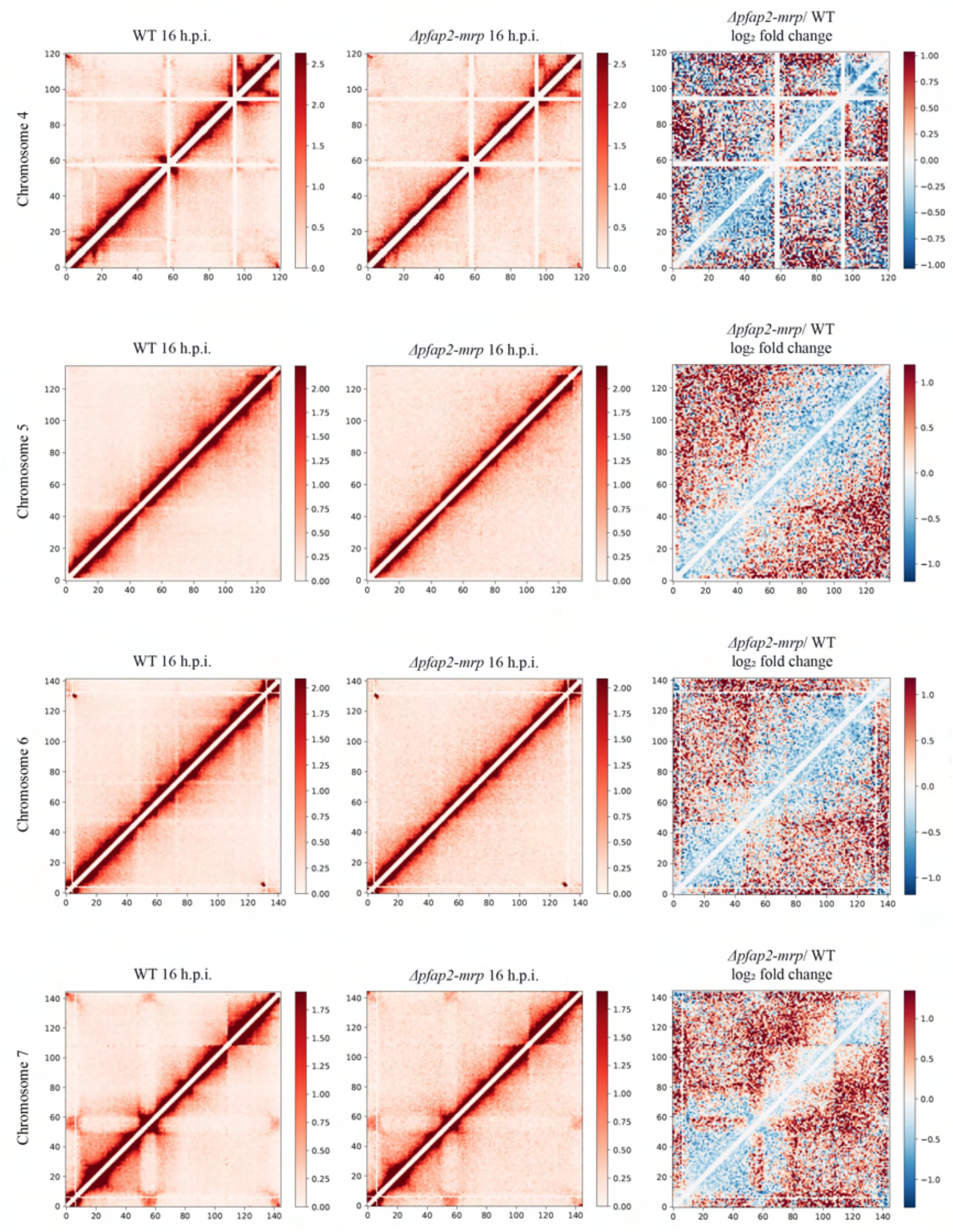
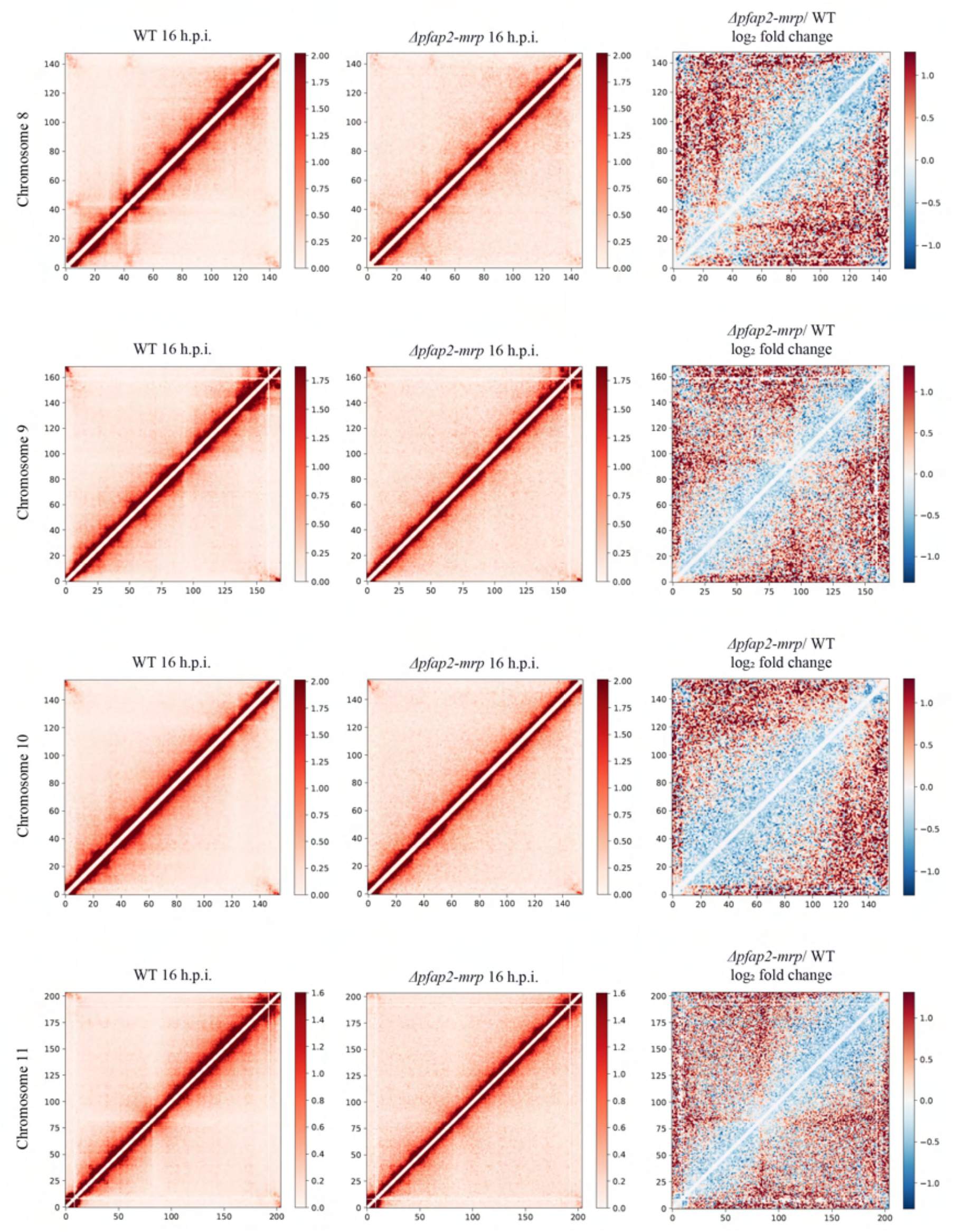
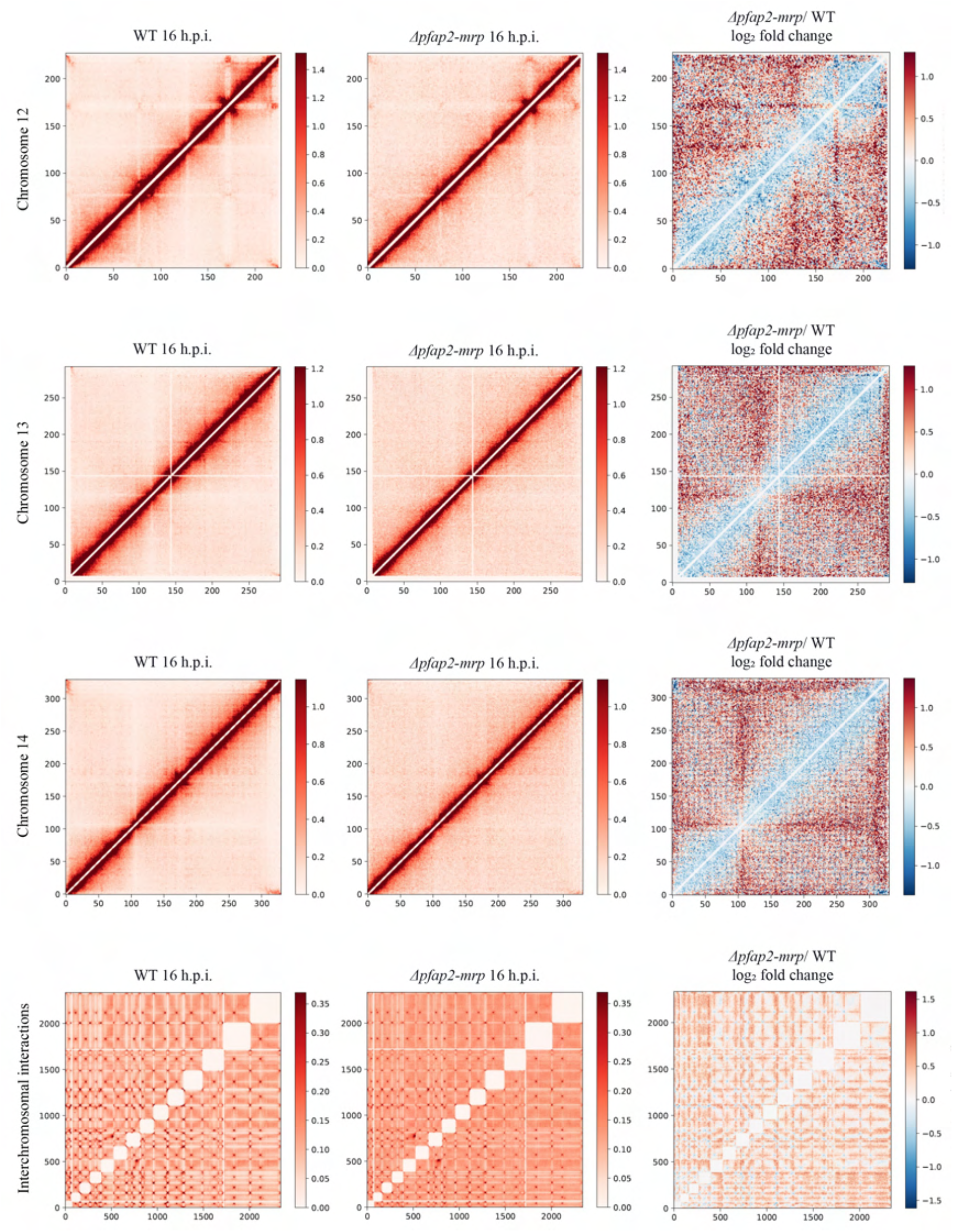
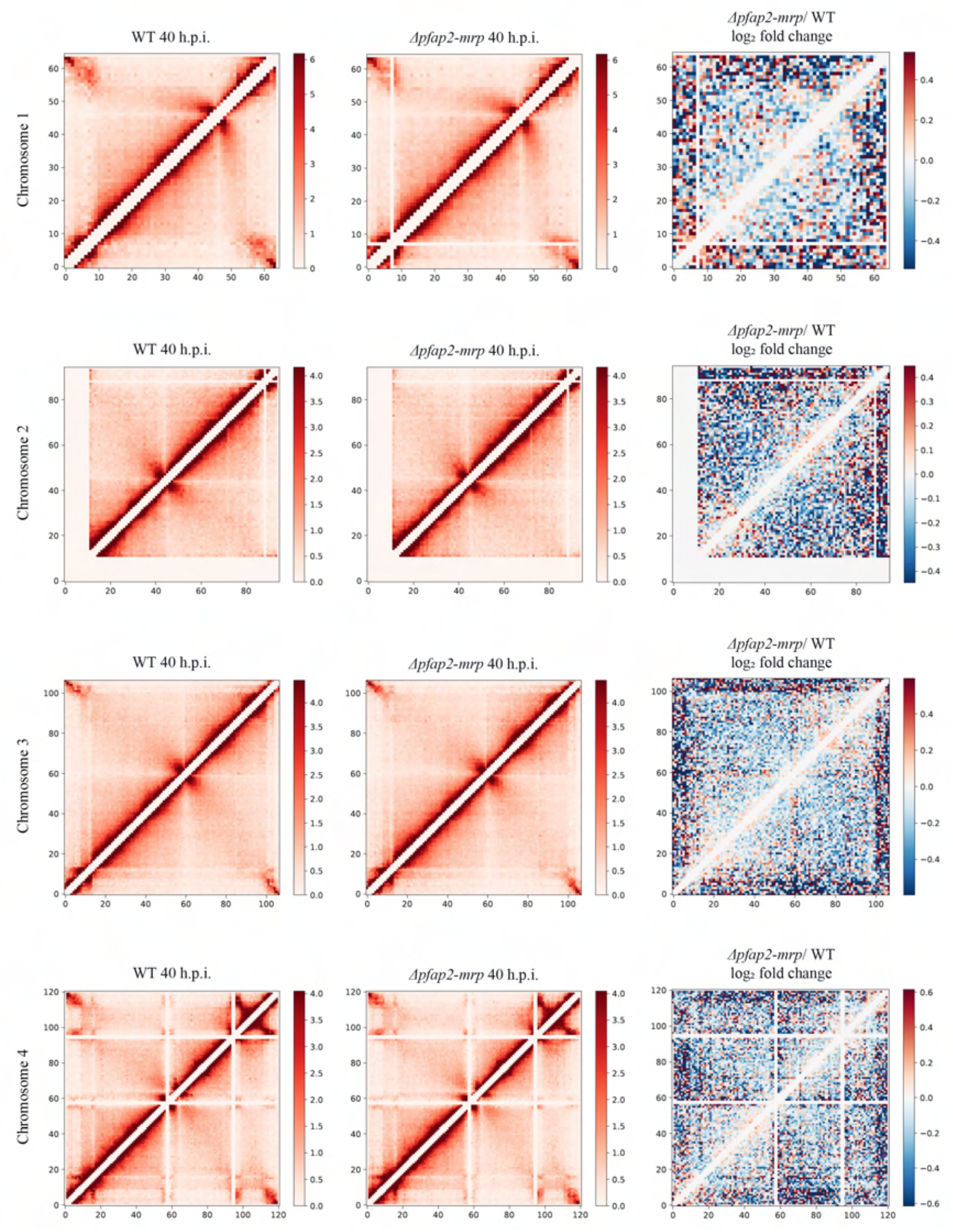
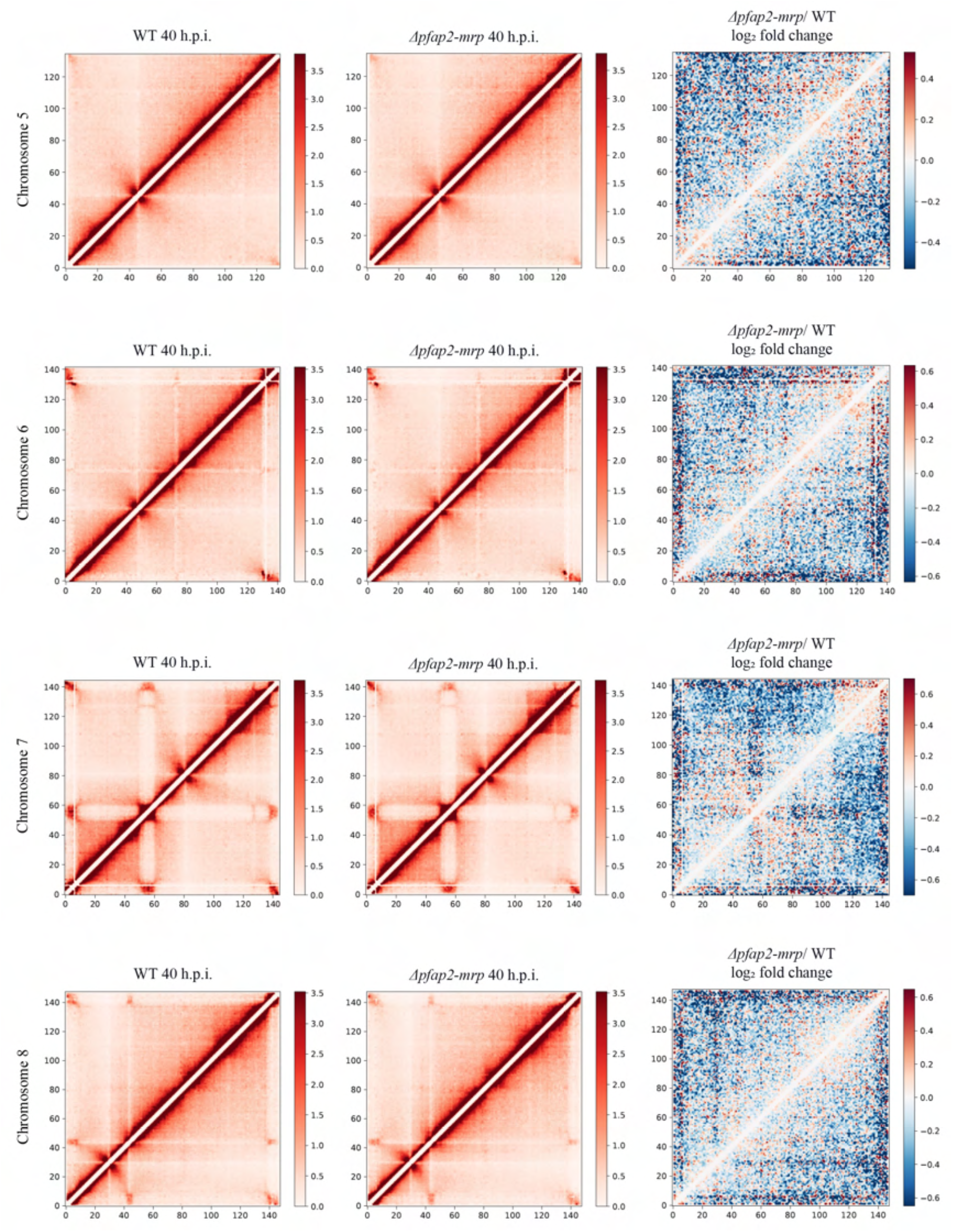
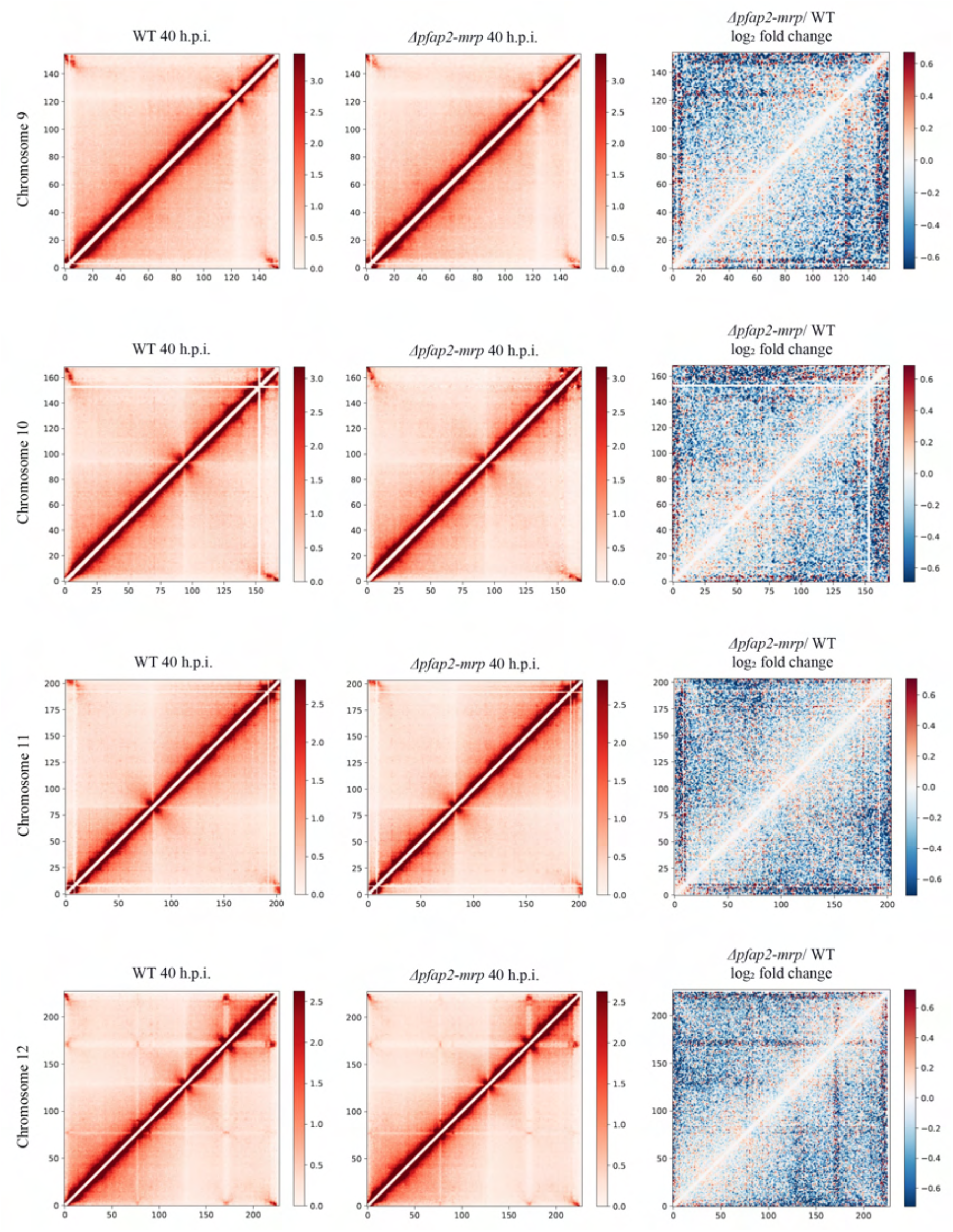
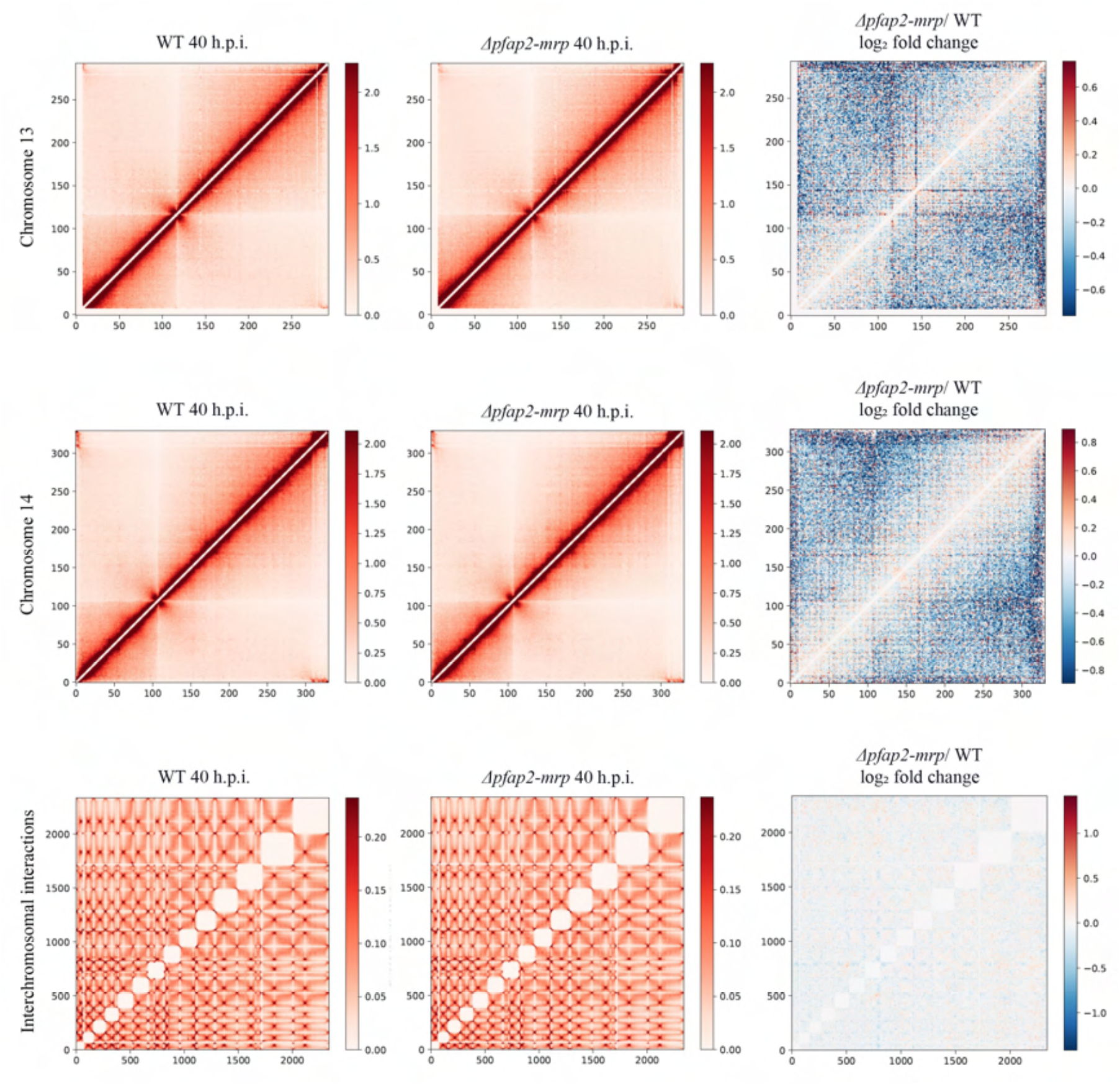
Visualizations of in-situ Hi-C generated chromatin interactions. ICE-normalized contact count heatmaps at 10kb resolution of intrachromosomal interactions for the 14 chromosomes, as well as a genome-wide contact count heatmap showing interchromosomal interactions, are given for both the 16 h.p.i. and 40 h.p.i. time points. Each row represents a single chromosome of the wild type (left), *Δpfap2-mrp* (center), and log₂ fold change differential interactions (right). The data for the two biological replicates were merged using a weighted average based on the total read count and then counts-per-million normalized prior to generating heatmaps. The scale of the legend was also normalized to improve comparisons between WT and *Δpfap2-mrp*. For each bin *i,* all interactions within *i* +/-2 are set to 0 (see white line at diagonal) to enhance visualization of remaining bins due to intra-bin and very short-range contacts being significantly higher

**Extended Data Fig. 9.**
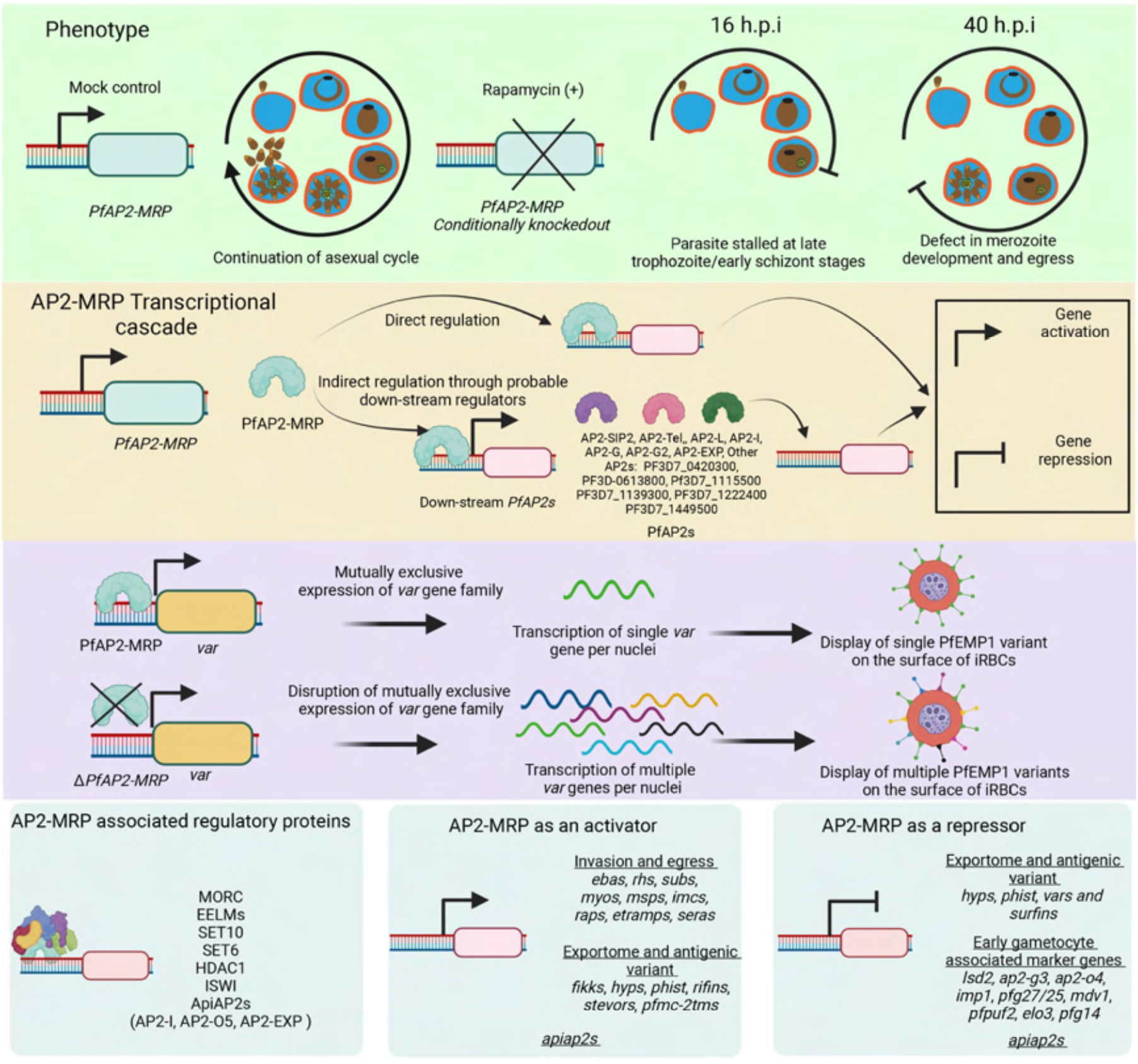
PfAP2-MRP is a master regulator of malaria pathogenesis. Deletion of *pfap2-mrp* before the first peak of expression at 16 h.p.i. affects parasite development beyond late trophozoite/early schizont stages. Deletion of *pfap2-mrp* well before 40 h.p.i. but after the first peak of expression at 16 h.p.i. affects merozoite development and blocks parasite egress from infected RBCs. At this late stage of intraerythrocytic development, the second peak of *pfap2-mrp* expression activates many genes associated with invasion, egress, antigenic variation, host cell remodeling and protein phosphorylation, either directly by binding to their promoter or indirectly through other downstream ApiAP2 transcription factors and regulators. PfAP2-MRP is a direct repressor of *var* genes and an indirect repressor of many gametocytogenesis-associated marker genes. Deletion of *pfap2-mrp* derepresses expression of most of *var* genes leading to the displayed of the corresponding PfEMP1 on the iRBC surface. PfAP2-MRP acts as a direct activator of many other genes such as *rifins*, *stevors*, and *pfmc-2tms* coding for antigenically variant proteins, and binds to the promoter of 14 other ApiAP2s (50% of all *P. falciparum* ApiAP2 genes), suggesting that it is an upstream regulator of gene expression cascades during the IDC. PfAP2-MRP associates with many known and putative histone modifiers and chromatin remodelers that probably participate in PfAP2-MRP associated gene regulation. Altogether, PfAP2-MRP regulates the expression of most known pathogenic factors (associated with antigenic variation and parasite growth) in *P. falciparum* suggesting it is a master regulator of malaria pathogenesis.

## References

1 Cowman, A. F., Tonkin, C. J., Tham, W. H. & Duraisingh, M. T. The Molecular Basis of Erythrocyte Invasion by Malaria Parasites. Cell Host Microbe 22, 232–245, doi:10.1016/j.chom.2017.07.003 (2017).

2 Boddey, J. A. & Cowman, A. F. Plasmodium nesting: remaking the erythrocyte from the inside out. Annu Rev Microbiol 67, 243–269, doi:10.1146/annurev-micro-092412-155730 (2013).

3 Tan, M. S. Y. & Blackman, M. J. Malaria parasite egress at a glance. J Cell Sci 134, doi:10.1242/jcs.257345 (2021).

4 Cortes, A. & Deitsch, K. W. Malaria Epigenetics. Cold Spring Harb Perspect Med 7, doi:10.1101/cshperspect.a025528 (2017).

5 Toenhake, C. G. et al. Chromatin Accessibility-Based Characterization of the Gene Regulatory Network Underlying Plasmodium falciparum Blood-Stage Development. Cell Host Microbe 23, 557–569 e559, doi:10.1016/j.chom.2018.03.007 (2018).

6 Iwanaga, S., Kaneko, I., Kato, T. & Yuda, M. Identification of an AP2-family protein that is critical for malaria liver stage development. PLoS One 7, e47557, doi:10.1371/journal.pone.0047557 (2012).

7 Kafsack, B. F. et al. A transcriptional switch underlies commitment to sexual development in malaria parasites. Nature 507, 248–252, doi:10.1038/nature12920 (2014).

8 Painter, H. J., Campbell, T. L. & Llinas, M. The Apicomplexan AP2 family: integral factors regulating Plasmodium development. Mol Biochem Parasitol 176, 1–7, doi:10.1016/j.molbiopara.2010.11.014 (2011).

9 Tinto-Font, E. et al. A heat-shock response regulated by the PfAP2-HS transcription factor protects human malaria parasites from febrile temperatures. Nat Microbiol 6, 1163–1174, doi:10.1038/s41564-021-00940-w (2021).

10 Yuda, M., Iwanaga, S., Shigenobu, S., Kato, T. & Kaneko, I. Transcription factor AP2-Sp and its target genes in malarial sporozoites. Mol Microbiol 75, 854–863, doi:10.1111/j.1365-2958.2009.07005.x (2010).

11 Yuda, M. et al. Identification of a transcription factor in the mosquito-invasive stage of malaria parasites. Mol Microbiol 71, 1402–1414, doi:10.1111/j.1365-2958.2009.06609.x (2009).

12 Collins, C. R. et al. Malaria parasite cGMP-dependent protein kinase regulates blood stage merozoite secretory organelle discharge and egress. PLoS Pathog 9, e1003344, doi:10.1371/journal.ppat.1003344 (2013).

13 Knuepfer, E., Napiorkowska, M., van Ooij, C. & Holder, A. A. Generating conditional gene knockouts in Plasmodium – a toolkit to produce stable DiCre recombinase-expressing parasite lines using CRISPR/Cas9. Sci Rep 7, 3881, doi:10.1038/s41598-017-03984-3 (2017).

14 Subudhi, A. K. et al. Malaria parasites regulate intra-erythrocytic development duration via serpentine receptor 10 to coordinate with host rhythms. Nat Commun 11, 2763, doi:10.1038/s41467-020-16593-y (2020).

15 Gomes, A. R. et al. A genome-scale vector resource enables high-throughput reverse genetic screening in a malaria parasite. Cell Host Microbe 17, 404–413, doi:10.1016/j.chom.2015.01.014 (2015).

16 Zhang, M. et al. Uncovering the essential genes of the human malaria parasite Plasmodium falciparum by saturation mutagenesis. Science 360, doi:10.1126/science.aap7847 (2018).

17 Thomas, J. A. et al. A protease cascade regulates release of the human malaria parasite Plasmodium falciparum from host red blood cells. Nat Microbiol 3, 447–455, doi:10.1038/s41564-018-0111-0 (2018).

18 Scherf, A., Lopez-Rubio, J. J. & Riviere, L. Antigenic variation in Plasmodium falciparum. Annu Rev Microbiol 62, 445–470, doi:10.1146/annurev.micro.61.080706.093134 (2008).

19 Baker, D. A. et al. Cyclic nucleotide signalling in malaria parasites. Open Biol 7, doi:10.1098/rsob.170213 (2017).

20 Singh, S. & Chitnis, C. E. Molecular Signaling Involved in Entry and Exit of Malaria Parasites from Host Erythrocytes. Cold Spring Harb Perspect Med 7, doi:10.1101/cshperspect.a026815 (2017).

21 Sargeant, T. J. et al. Lineage-specific expansion of proteins exported to erythrocytes in malaria parasites. Genome Biol 7, R12, doi:10.1186/gb-2006-7-2-r12 (2006).

22 Siddiqui, G., Proellochs, N. I. & Cooke, B. M. Identification of essential exported Plasmodium falciparum protein kinases in malaria-infected red blood cells. Br J Haematol 188, 774–783, doi:10.1111/bjh.16219 (2020).

23 Howick, V. M. et al. The Malaria Cell Atlas: Single parasite transcriptomes across the complete Plasmodium life cycle. Science 365, doi:10.1126/science.aaw2619 (2019).

24 Taylor, T. E. et al. Intravenous immunoglobulin in the treatment of paediatric cerebral malaria. Clin Exp Immunol 90, 357–362, doi:10.1111/j.1365-2249.1992.tb05851.x (1992).

25 Chan, J. A., Fowkes, F. J. & Beeson, J. G. Surface antigens of Plasmodium falciparum-infected erythrocytes as immune targets and malaria vaccine candidates. Cell Mol Life Sci 71, 3633–3657, doi:10.1007/s00018-014-1614-3 (2014).

26 Gulati, S. et al. Profiling the Essential Nature of Lipid Metabolism in Asexual Blood and Gametocyte Stages of Plasmodium falciparum. Cell Host Microbe 18, 371–381, doi:10.1016/j.chom.2015.08.003 (2015).

27 Josling, G. A. & Llinas, M. Sexual development in Plasmodium parasites: knowing when it’s time to commit. Nat Rev Microbiol 13, 573–587, doi:10.1038/nrmicro3519 (2015).

28 Poran, A. et al. Single-cell RNA sequencing reveals a signature of sexual commitment in malaria parasites. Nature 551, 95–99, doi:10.1038/nature24280 (2017).

29 Shang, X. et al. Genome-wide landscape of ApiAP2 transcription factors reveals a heterochromatin-associated regulatory network during Plasmodium falciparum blood-stage development. Nucleic Acids Res, doi:10.1093/nar/gkac176 (2022).

30 Santos, J. M. et al. Red Blood Cell Invasion by the Malaria Parasite Is Coordinated by the PfAP2-I Transcription Factor. Cell Host Microbe 21, 731–741 e710, doi:10.1016/j.chom.2017.05.006 (2017).

31 Josling, G. A. et al. Dissecting the role of PfAP2-G in malaria gametocytogenesis. Nat Commun 11, 1503, doi:10.1038/s41467-020-15026-0 (2020).

32 Volz, J. C. et al. PfSET10, a Plasmodium falciparum methyltransferase, maintains the active var gene in a poised state during parasite division. Cell Host Microbe 11, 7–18, doi:10.1016/j.chom.2011.11.011 (2012).

33 Volz, J. et al. Potential epigenetic regulatory proteins localise to distinct nuclear sub-compartments in Plasmodium falciparum. Int J Parasitol 40, 109–121, doi:10.1016/j.ijpara.2009.09.002 (2010).

34 Bryant, J. M. et al. Exploring the virulence gene interactome with CRISPR/dCas9 in the human malaria parasite. Mol Syst Biol 16, e9569, doi:10.15252/msb.20209569 (2020).

35 Farhat, D. C. et al. A MORC-driven transcriptional switch controls Toxoplasma developmental trajectories and sexual commitment. Nat Microbiol 5, 570–583, doi:10.1038/s41564-020-0674-4 (2020).

36 Andrews, K. T., Haque, A. & Jones, M. K. HDAC inhibitors in parasitic diseases. Immunol Cell Biol 90, 66–77, doi:10.1038/icb.2011.97 (2012).

37 Ay, F. et al. Three-dimensional modeling of the P. falciparum genome during the erythrocytic cycle reveals a strong connection between genome architecture and gene expression. Genome Res 24, 974–988, doi:10.1101/gr.169417.113 (2014).

38 Bunnik, E. M. et al. Changes in genome organization of parasite-specific gene families during the Plasmodium transmission stages. Nat Commun 9, 1910, doi:10.1038/s41467-018-04295-5 (2018).

39 Bunnik, E. M. et al. Comparative 3D genome organization in apicomplexan parasites. Proc Natl Acad Sci U S A 116, 3183–3192, doi:10.1073/pnas.1810815116 (2019).

40 Oliver, S. Guilt-by-association goes global. Nature 403, 601–603, doi:10.1038/35001165 (2000).

41 Nagaoka, H. et al. PfMSA180 is a novel Plasmodium falciparum vaccine antigen that interacts with human erythrocyte integrin associated protein (CD47). Sci Rep 9, 5923, doi:10.1038/s41598-019-42366-9(2019).

42 Liffner, B. et al. PfCERLI1 is a conserved rhoptry associated protein essential for Plasmodium falciparum merozoite invasion of erythrocytes. Nat Commun 11, 1411, doi:10.1038/s41467-020-15127-w (2020).

43 Wichers, J. S. et al. Identification of novel inner membrane complex and apical annuli proteins of the malaria parasite Plasmodium falciparum. Cell Microbiol 23, e13341, doi:10.1111/cmi.13341 (2021).

44 Tarr, S. J. et al. A malaria parasite subtilisin propeptide-like protein is a potent inhibitor of the egress protease SUB1. Biochem J 477, 525–540, doi:10.1042/BCJ20190918 (2020).

45 Moon, R. W. et al. Adaptation of the genetically tractable malaria pathogen Plasmodium knowlesi to continuous culture in human erythrocytes. Proc Natl Acad Sci U S A 110, 531–536, doi:10.1073/pnas.1216457110 (2013).

46 Jones, M. L. et al. A versatile strategy for rapid conditional genome engineering using loxP sites in a small synthetic intron in Plasmodium falciparum. Sci Rep 6, 21800, doi:10.1038/srep21800 (2016).

47 Bolger, A. M., Lohse, M. & Usadel, B. Trimmomatic: a flexible trimmer for Illumina sequence data. Bioinformatics 30, 2114–2120, doi:10.1093/bioinformatics/btu170 (2014).

48 Kim, D., Langmead, B. & Salzberg, S. L. HISAT: a fast spliced aligner with low memory requirements. Nat Methods 12, 357–360, doi:10.1038/nmeth.3317 (2015).

49 Liao, Y., Smyth, G. K. & Shi, W. featureCounts: an efficient general purpose program for assigning sequence reads to genomic features. Bioinformatics 30, 923–930, doi:10.1093/bioinformatics/btt656 (2014).

50 McCarthy, D. J., Chen, Y. & Smyth, G. K. Differential expression analysis of multifactor RNA-Seq experiments with respect to biological variation. Nucleic Acids Res 40, 4288–4297, doi:10.1093/nar/gks042 (2012).

51 Ritchie, M. E. et al. limma powers differential expression analyses for RNA-sequencing and microarray studies. Nucleic Acids Res 43, e47, doi:10.1093/nar/gkv007 (2015).

52 Love, M. I., Huber, W. & Anders, S. Moderated estimation of fold change and dispersion for RNA-seq data with DESeq2. Genome Biol 15, 550, doi:10.1186/s13059-014-0550-8 (2014).

53 Pfaffl, M. W. A new mathematical model for relative quantification in real-time RT-PCR. Nucleic Acids Res 29, e45, doi:10.1093/nar/29.9.e45 (2001).

54 Jiang, L. et al. PfSETvs methylation of histone H3K36 represses virulence genes in Plasmodium falciparum. Nature 499, 223–227, doi:10.1038/nature12361 (2013).

55 Zeeshan, M. et al. Real-time dynamics of Plasmodium NDC80 reveals unusual modes of chromosome segregation during parasite proliferation. J Cell Sci 134, doi:10.1242/jcs.245753 (2020).

56 Li, H. et al. The Sequence Alignment/Map format and SAMtools. Bioinformatics 25, 2078–2079, doi:10.1093/bioinformatics/btp352 (2009).

57 Ramirez, F., Dundar, F., Diehl, S., Gruning, B. A. & Manke, T. deepTools: a flexible platform for exploring deep-sequencing data. Nucleic Acids Res 42, W187–191, doi:10.1093/nar/gku365 (2014).

58 Thorvaldsdottir, H., Robinson, J. T. & Mesirov, J. P. Integrative Genomics Viewer (IGV): high-performance genomics data visualization and exploration. Brief Bioinform 14, 178–192, doi:10.1093/bib/bbs017 (2013).

59 Zhang, Y. et al. Model-based analysis of ChIP-Seq (MACS). Genome Biol 9, R137, doi:10.1186/gb-2008-9-9-r137 (2008).

60 Quinlan, A. R. & Hall, I. M. BEDTools: a flexible suite of utilities for comparing genomic features. Bioinformatics 26, 841–842, doi:10.1093/bioinformatics/btq033 (2010).

61 Lun, A. T. L. et al. EmptyDrops: distinguishing cells from empty droplets in droplet-based single-cell RNA sequencing data. Genome Biol 20, 63, doi:10.1186/s13059-019-1662-y (2019).

62 McCarthy, D. J., Campbell, K. R., Lun, A. T. & Wills, Q. F. Scater: pre-processing, quality control, normalization and visualization of single-cell RNA-seq data in R. Bioinformatics 33, 1179–1186, doi:10.1093/bioinformatics/btw777 (2017).

63 Lun, A. T., McCarthy, D. J. & Marioni, J. C. A step-by-step workflow for low-level analysis of single-cell RNA-seq data with Bioconductor. F1000Res 5, 2122, doi: 10.12688/f1000research.9501.2 (2016).

64 Hao, Y. et al. Integrated analysis of multimodal single-cell data. Cell 184, 3573–3587 e3529, doi:10.1016/j.cell.2021.04.048 (2021).

65 Real, E. et al. A single-cell atlas of Plasmodium falciparum transmission through the mosquito. Nat Commun 12, 3196, doi:10.1038/s41467-021-23434-z (2021).

66 Aran, D. et al. Reference-based analysis of lung single-cell sequencing reveals a transitional profibrotic macrophage. Nat Immunol 20, 163–172, doi:10.1038/s41590-018-0276-y (2019).

67 de Kanter, J. K., Lijnzaad, P., Candelli, T., Margaritis, T. & Holstege, F. C. P. CHETAH: a selective, hierarchical cell type identification method for single-cell RNA sequencing. Nucleic Acids Res 47, e95, doi:10.1093/nar/gkz543 (2019).

68 Bailey, T. L. DREME: motif discovery in transcription factor ChIP-seq data. Bioinformatics 27, 1653–1659, doi:10.1093/bioinformatics/btr261 (2011).

69 Gupta, S., Stamatoyannopoulos, J. A., Bailey, T. L. & Noble, W. S. Quantifying similarity between motifs. Genome Biol 8, R24, doi:10.1186/gb-2007-8-2-r24 (2007).

70 Campbell, T. L., De Silva, E. K., Olszewski, K. L., Elemento, O. & Llinas, M. Identification and genome-wide prediction of DNA binding specificities for the ApiAP2 family of regulators from the malaria parasite. PLoS Pathog 6, e1001165, doi:10.1371/journal.ppat.1001165 (2010).

71 Rao, S. S. et al. A 3D map of the human genome at kilobase resolution reveals principles of chromatin looping. Cell 159, 1665–1680, doi:10.1016/j.cell.2014.11.021 (2014).

72 Gupta, M. K., Lenz, T. & Le Roch, K. G. Chromosomes Conformation Capture Coupled with Next-Generation Sequencing (Hi-C) in Plasmodium falciparum. Methods Mol Biol 2369, 15–25, doi:10.1007/978-1-0716-1681-9_2 (2021).

73 Varoquaux, N., Ay, F., Noble, W. S. & Vert, J. P. A statistical approach for inferring the 3D structure of the genome. Bioinformatics 30, i26–33, doi:10.1093/bioinformatics/btu268 (2014).

74 Goddard, T. D., et al. UCSF ChimeraX: Meeting modern challenges in visualization and analysis. Protein Sci 27, 14–25, doi:10.1002/pro.3235 (2018).

