## Supplementary Information for "PfAP2-MRP DNA-binding protein is a master regulator of parasite pathogenesis during malaria parasite blood stages"

**This file contains supplementary discussion, references and legends for supplementary data 1-6.**

**Supplementary Discussion**

In a previous study, it was demonstrated that PfAP2-I and PfAP2-G bind to the same promoter regions of many invasion-associated genes^1^. Our ChIP-seq data and motif enrichment analysis indicate that PfAP2-MRP binds to the same regions of many of these genes, suggesting a complex gene regulatory system, important for both asexual parasite growth and sexual commitment. For example, a combinatorial binding of transcription factors to the same promoter has been proposed as a mechanism to increase specificity and facilitate fine-tuning^2^. The binding of PfAP2-MRP and other PfAP2 transcription factors to the same promoter regions is consistent with such a mechanism. A recent study^3^, has identified that ApiAP2s are regulated by multiple other ApiAP2s. A couple of such cases are *pfap2-hc* and PF3D7_0613800 (another uncharacterized ApiAP2), which are regulated by at least 8 ApiAP2s including themselves, which further supports the role of combinatorial based gene regulation in malaria parasites.

The gene transcriptional activation and suppression of PfAP2-MRP, suggest that PfAP2-MRP may have a role in epigenetic regulation of gene expression, acting together with other factors. Known and putative histone modifiers and chromatin remodelers were identified in complex with PfAP2-MRP by IP and mass spectrometry, supporting the idea that PfAP2-MRP recruits these epigenetic regulators. Recently, PfAP2-MRP was identified as a protein associated with *var* gene promoter regions together with chromatin remodelers enriched at the same location^4^, consistent with our data. We identified PfEELM2 and PfMORC associated PfAP2-MRP at both 16 and 40 h.p.i., consistent with a *P. berghei* study in which PBANKA_0939100, the ortholog of PfAP2-MRP, was shown to interact with PbEELM2 (PBANKA_234600) and PbMORC (PBANKA_1331400), in schizonts (Hillier et al., 2019). EELM2 and MORC proteins are often associated with histone deacetylase in a complex with chromatin remodeling activities associated with gene suppression^5^. Recently *Toxoplasma gondii* MORC was shown to be a transcriptional repressor of sexual commitment when interacting with AP2 DNA-binding proteins^6^. In previous work^7^, we showed that PfMORC has two expression peaks in the IDC, coincident with those of PfAP2-MRP (Pearson correlation 0.72). It is therefore possible that PfAP2-MRP and PfMORC interact together to repress *var* gene expression. As we also identified known transcriptional activators such as PfSET10, PfSET6, and PfISWI interacting with PfAP2-MRP, we propose that PfAP2-MRP may recruit histone modifiers and chromatin remodelers to alter the cis-chromatin structure, leading to either gene activation or repression. This would be consistent with roles for PfAP2-MRP as both activator (of some genes associated with antigenic variation, host cell modification, egress and invasion) and repressor (of *var* and gametocytogenesis-associated genes).

Signal transduction pathways that rely on protein kinases and phosphatases have crucial roles in the parasite life cycle^8,9^ including the merozoite egress and erythrocyte invasion stages where there are substantial differences in the phosphoproteomes of intracellular schizonts and extracellular merozoites^10^. Many of the proteins phosphorylated in merozoites probably participate in egress, movement, and invasion, and are phosphorylated by kinases such as PKG, PKA, CDPK1, and CDPK5. Genes encoding these kinases were down-regulated in *Δpfap2-mrp* parasites at 40 h.p.i., suggesting another significant role for PfAP2-MRP in mechanisms regulating these pathogenic processes.

FIKK kinases are orphan kinases restricted to apicomplexan parasites. A single FIKK gene is present in all *Plasmodium* genomes. However, in species of the Laverania clade, the family has been expanded to between 18 and 26 distinct members, with 21 genes in the *fikk* family of *P. falciparu* ^11,12^. Eighteen of these genes code for proteins with a putative signal sequence, and suggested to be exported into the iRBC to mediate the parasite-induced modification, which is central to pathogenesis^13^. Other roles are also possible, for example it was shown that FIKK3 co-localizes with PfRAMA in the rhoptry bulb^14^. A total of 12 *fikk* genes, including *fikk3,* were identified as down-regulated in *Δpfap2-mrp* parasites at 16 and 40 h.p.i., consistent with a role in host cell remodeling and invasion.

Increased expression of many known and putative gametocyte-marker genes was observed in *Δpfap2-mrp* parasites at both 16 and 40 h.p.i., including genes encoding recently identified putative transcriptional regulators of gametocytogenesis such as lysine-specific demethylase (LSD2, a putative histone demethylase), AP2-O4, AP2-G3, and AP2 (PF3D7_1139300)^15^. LSD2 and AP2 were identified as potential regulators driving the expression of genes for gametocyte development in committed schizonts^16^. AP2-G3, strongly up-regulated in *Δpfap2-mrp* parasites, probably plays an essential role in gametocyte production as a regulator upstream of AP2-G^17^. Other up-regulated genes include those for male gametocyte development (PfMDV-1) and an mRNA binding protein (PfPuf2) important in male and female gametocyte development, respectively^18,19^. These data suggest that PfAP2-MRP inhibits commitment to sexual stage development acting through an indirect regulator as we did not observe binding of PfAP2-MRP to the promoters of these genes.

**Supplementary Data 1** | Differentially expressed genes at 16 h.p.i. and 40 h.p.i. in *Δpfap2-mrp* parasites

List of differentially expressed genes at 16 h.p.i. and 40 h.p.i. after disruption of first peak and second peak of *pfap2-mrp* expression, respectively.

**Supplementary Data 2** | Gene ontology enrichment analysis of differentially expressed genes at 16 h.p.i. and 40 h.p.i. in *Δpfap2-mrp* parasites

**Supplementary Data 3** | ChIP-seq peaks identified at 16 h.p.i. and 40 h.p.i.

**Supplementary Data 4** | Gene ontology enrichment analysis of genes whose promoter regions or the gene body were bound by PfAP2-MRP at 16 h.p.i. and 40 h.p.i.

**Supplementary Data 5** | PfAP2-MRP associated proteins identified at 16 and 40 h.p.i. using ChIP followed by Mass Spectrometry

**Supplementary Data 6** | Oligos used in this study

**References**

1	Josling, G. A. *et al.* Dissecting the role of PfAP2-G in malaria gametocytogenesis. *Nat Commun* **11**, 1503, doi:10.1038/s41467-020-15026-0 (2020).

2	Reiter, F., Wienerroither, S. & Stark, A. Combinatorial function of transcription factors and cofactors. *Curr Opin Genet Dev* **43**, 73-81, doi:10.1016/j.gde.2016.12.007 (2017).

3	Shang, X. *et al.* A cascade of transcriptional repression determines sexual commitment and development in Plasmodium falciparum. *Nucleic Acids Res* **49**, 9264-9279, doi:10.1093/nar/gkab683 (2021).

4	Bryant, J. M. *et al.* Exploring the virulence gene interactome with CRISPR/dCas9 in the human malaria parasite. *Mol Syst Biol* **16**, e9569, doi:10.15252/msb.20209569 (2020).

5	Solari, F., Bateman, A. & Ahringer, J. The Caenorhabditis elegans genes egl-27 and egr-1 are similar to MTA1, a member of a chromatin regulatory complex, and are redundantly required for embryonic patterning. *Development* **126**, 2483-2494 (1999).

6	Farhat, D. C. *et al.* A MORC-driven transcriptional switch controls Toxoplasma developmental trajectories and sexual commitment. *Nat Microbiol* **5**, 570-583, doi:10.1038/s41564-020-0674-4 (2020).

7	Subudhi, A. K. *et al.* Malaria parasites regulate intra-erythrocytic development duration via serpentine receptor 10 to coordinate with host rhythms. *Nat Commun* **11**, 2763, doi:10.1038/s41467-020-16593-y (2020).

8	Baker, D. A. *et al.* Cyclic nucleotide signalling in malaria parasites. *Open Biol* **7**, doi:10.1098/rsob.170213 (2017).

9	Singh, S. & Chitnis, C. E. Molecular Signaling Involved in Entry and Exit of Malaria Parasites from Host Erythrocytes. *Cold Spring Harb Perspect Med* **7**, doi:10.1101/cshperspect.a026815 (2017).

10	Lasonder, E., Green, J. L., Grainger, M., Langsley, G. & Holder, A. A. Extensive differential protein phosphorylation as intraerythrocytic Plasmodium falciparum schizonts develop into extracellular invasive merozoites. *Proteomics* **15**, 2716-2729, doi:10.1002/pmic.201400508 (2015).

11	Proellocks, N. I., Coppel, R. L., Mohandas, N. & Cooke, B. M. Malaria Parasite Proteins and Their Role in Alteration of the Structure and Function of Red Blood Cells. *Adv Parasitol* **91**, 1-86, doi:10.1016/bs.apar.2015.09.002 (2016).

12	Ward, P., Equinet, L., Packer, J. & Doerig, C. Protein kinases of the human malaria parasite Plasmodium falciparum: the kinome of a divergent eukaryote. *BMC Genomics* **5**, 79, doi:10.1186/1471-2164-5-79 (2004).

13	Davies, H. *et al.* An exported kinase family mediates species-specific erythrocyte remodelling and virulence in human malaria. *Nat Microbiol* **5**, 848-863, doi:10.1038/s41564-020-0702-4 (2020).

14	Siddiqui, G., Proellochs, N. I. & Cooke, B. M. Identification of essential exported Plasmodium falciparum protein kinases in malaria-infected red blood cells. *Br J Haematol* **188**, 774-783, doi:10.1111/bjh.16219 (2020).

15	Zhang, M. *et al.* Uncovering the essential genes of the human malaria parasite Plasmodium falciparum by saturation mutagenesis. *Science* **360**, doi:10.1126/science.aap7847 (2018).

16	Poran, A. *et al.* Single-cell RNA sequencing reveals a signature of sexual commitment in malaria parasites. *Nature* **551**, 95-99, doi:10.1038/nature24280 (2017).

17	Josling, G. A. & Llinas, M. Sexual development in Plasmodium parasites: knowing when it's time to commit. *Nat Rev Microbiol* **13**, 573-587, doi:10.1038/nrmicro3519 (2015).

18	Furuya, T. *et al.* Disruption of a Plasmodium falciparum gene linked to male sexual development causes early arrest in gametocytogenesis. *Proc Natl Acad Sci U S A* **102**, 16813-16818, doi:10.1073/pnas.0501858102 (2005).

19	Miao, J. *et al.* The Puf-family RNA-binding protein PfPuf2 regulates sexual development and sex differentiation in the malaria parasite Plasmodium falciparum. *J Cell Sci* **123**, 1039-1049, doi:10.1242/jcs.059824 (2010).
